# Structures of Epstein-Barr virus and Kaposi’s sarcoma-associated herpesvirus virions reveal species-specific tegument and envelope features

**DOI:** 10.1101/2024.07.09.602672

**Authors:** James Zhen, Jia Chen, Haigen Huang, Shiqing Liao, Shiheng Liu, Yan Yuan, Ren Sun, Richard Longnecker, Ting-Ting Wu, Z. Hong Zhou

**Affiliations:** Department of Microbiology, Immunology, and Molecular Genetics, University of California, Los Angeles (UCLA), Los Angeles, CA, USA; California NanoSystems Institute, UCLA, Los Angeles, CA, USA; Molecular Biology Institute, UCLA, Los Angeles, CA, USA; Department of Microbiology and Immunology, Feinberg School of Medicine, Northwestern University, Chicago, IL, USA; Department of Molecular and Medical Pharmacology, David Geffen School of Medicine, UCLA, Los Angeles, CA, USA; Department of Microbiology, School of Dental Medicine, University of Pennsylvania, Philadelphia, PA, USA

## Abstract

Epstein-Barr virus (EBV) and Kaposi’s sarcoma-associated herpesvirus (KSHV) are classified into the gammaherpesvirus subfamily of *Herpesviridae*, which stands out from its alpha- and betaherpesvirus relatives due to the tumorigenicity of its members. Although structures of human alpha- and betaherpesviruses by cryogenic electron tomography (cryoET) have been reported, reconstructions of intact human gammaherpesvirus virions remain elusive. Here, we structurally characterize extracellular virions of EBV and KSHV by deep learning-enhanced cryoET, resolving both previously known monomorphic capsid structures and previously unknown pleomorphic features beyond the capsid. Through subtomogram averaging and subsequent tomogram-guided sub-particle reconstruction, we determined the orientation of KSHV nucleocapsids from mature virions with respect to the portal to provide spatial context for the tegument within the virion. Both EBV and KSHV have an eccentric capsid position and polarized distribution of tegument. Tegument species span from the capsid to the envelope and may serve as scaffolds for tegumentation and envelopment. The envelopes of EBV and KSHV are less densely populated with glycoproteins than those of herpes simplex virus 1 and human cytomegalovirus, representative members of alpha- and betaherpesviruses, respectively. This population density of glycoproteins correlates with their relative infectivity against HEK293T cells. Also, we observed fusion protein gB trimers exist within triplet arrangements in addition to standalone complexes, which is relevant to understanding dynamic processes such as fusion pore formation. Taken together, this study reveals nuanced yet important differences in the tegument and envelope architectures among human herpesviruses and provides insights into their varied cell tropism and infection.

**Importance:** Discovered in 1964, Epstein-Barr virus (EBV) is the first identified human oncogenic virus and the founding member of the gammaherpesvirus subfamily. In 1994, another cancer-causing virus was discovered in lesions of AIDS patients and later named Kaposi’s sarcoma-associated herpesvirus (KSHV), the second human gammaherpesvirus. Despite the historical importance of EBV and KSHV, technical difficulties with isolating large quantities of these viruses and the pleiomorphic nature of their envelope and tegument layers have limited structural characterization of their virions. In this study, we employed the latest technologies in cryogenic electron microscopy (cryoEM) and tomography (cryoET) supplemented with an artificial intelligence-powered data processing software package to reconstruct 3D structures of the EBV and KSHV virions. We uncovered unique properties of the envelope glycoproteins and tegument layers of both EBV and KSHV. Comparison of these features with their non-tumorigenic counterparts provides insights into their relevance during infection.

## Introduction

Epstein-Barr virus (EBV) and Kaposi’s sarcoma-associated herpesvirus (KSHV) are two of the seven currently known human herpesviruses and comprise the gammaherpesvirus subfamily (1, 2). Like other herpesviruses, EBV and KSHV establish lifelong latent infections. However, unlike other herpesviruses, these gammaherpesvirus infections may manifest as cancers (1, 2). EBV is associated with Hodgkin’s lymphoma, Burkett’s lymphomas, and nasopharyngeal carcinomas (1, 2). Additionally, EBV has recently been identified as a causative agent of multiple sclerosis (3). KSHV is the etiological agent of Kaposi’s sarcoma, a highly prevalent cancer in AIDS patients (4, 5), and is the cause of other lymphoproliferative disorders, including primary effusion lymphoma and multicentric Castleman disease (6, 7). Structural determination of EBV and KSHV would improve our understanding of their pathogeneses and inform therapies to relieve their cancer burden.

The alpha-, beta-, and gammaherpesvirus subfamilies are related primarily by their morphological similarity (8). All herpesviruses are composed of a double-stranded DNA (dsDNA) genome within an icosahedral capsid that is surrounded by tegument and further enclosed by an envelope. Despite their shared overall morphology, herpesviruses are diverse in their host tropisms and pathologies. Gammaherpesviruses establish latency in B cells, whereas alpha- and betaherpesviruses establish latency in neurons and mononuclear cells, respectively (9). To facilitate this diversity, herpesviruses possess additional proteins unique to their individual species, particularly in the tegument and on the envelope (10). The envelope proteins and tegument proteins are crucial for host entry, cell tropism, immune evasion, and conditioning of the cellular environment (11–14). The protein differences between herpesviruses should also be reflected in their structural morphologies.

Early structural studies of herpesvirus structures revealed the architecture and molecular composition of the capsid (15–21). Advances in single particle cryogenic electron microscopy (cryoEM) during the past decade have since enabled structural determination of herpesvirus capsids and capsid-associated tegument components, including those of EBV and KSHV, at near-atomic resolution (22–38). However, individual intact virions are not well characterized due to the pleiomorphic nature of both their envelope and tegument, which prevents averaging. Cryogenic electron tomography (cryoET) generates 3D reconstructions of individual intact virions and enables observation and description of structural elements on the pleiomorphic envelope and within the tegument. For example, cryoET studies of intact virions have been performed for human alpha- and betaherpesvirus (39, 40). Tomograms of herpes simplex virus type 1 (HSV-1) revealed a polar tegument arrangement and glycoprotein distribution around an eccentric nucleocapsid (39). CryoET has also been used to characterize the fusion protein gB in its prefusion conformation on human cytomegalovirus (HCMV) virions (40). Recent developments in cryoET data processing, such as deep learning-based missing wedge correction (41), have increased interpretability of tomograms, including direct identification of HCMV portal complexes in enveloped A- and B-capsids (36).

Despite these advances, structural characterizations of human gammaherpesvirus virions in intact states are not available due to the difficulty of culturing them at high concentration. To date, only murine gammaherpesvirus-68 (MHV-68) virions and the naked capsids of KSHV have been interrogated (42, 43). Although EBV virions from the B95-8 system are still difficult to obtain, the development of the inducible iSLK cell line has improved KSHV production (44). This advancement, combined with recent developments in cryoET software (41, 45), enables an architectural description of human gammaherpesviruses that provide new insights into their assembly and immunogenicity.

In this study, we report structures and observations of intact EBV and KSHV determined by deep learning-enhanced cryoET. Using subtomogram averaging and tomogram-guided sub-particle reconstruction, we identify the locations of the EBV and KSHV capsid vertices and the KSHV portal to enable orientational contextualization of the tegument with respect to both the capsid and the envelope. Within the tegument, we observe thin strands along the envelope and between the capsid and envelope, which may have structural or scaffolding roles. Gammaherpesviruses have two- to threefold fewer presenting proteins on the envelope surface compared to HSV-1 and HCMV, which is correlated with their relative infectivity. The fusion protein gB is found in triplet clusters, and this arrangement may further inform the mechanism of host membrane fusion.

## Results

### Architecture of EBV and KSHV virions

We isolated EBV virions by sucrose gradient centrifugation from infectious media collected from B95-8 cells. Through cryoEM imaging of the sample, we observe scant EBV virions and an abundance of vesicles (Fig. 1a). Some EBV virions deviate from their common spherical depiction, but the presence of spherical vesicles co-isolated with them suggests that such deviations are not solely attributed to centrifugation. Gammaherpesvirus infection produces both virions and virus-like vesicles (46), the latter of which obstructs data collection and quality by increasing sample thickness and obscuring virions. Nonetheless, we proceeded to collect tilt series of this suboptimal EBV sample for the purpose of high-resolution reconstruction (Fig. 1b). In the low-dose images of EBV virion, we observe the leaflets of the bilayer envelope and the contours of the dsDNA genome within the capsid, which indicate that moderate resolution features are present.

**Figure 1.**
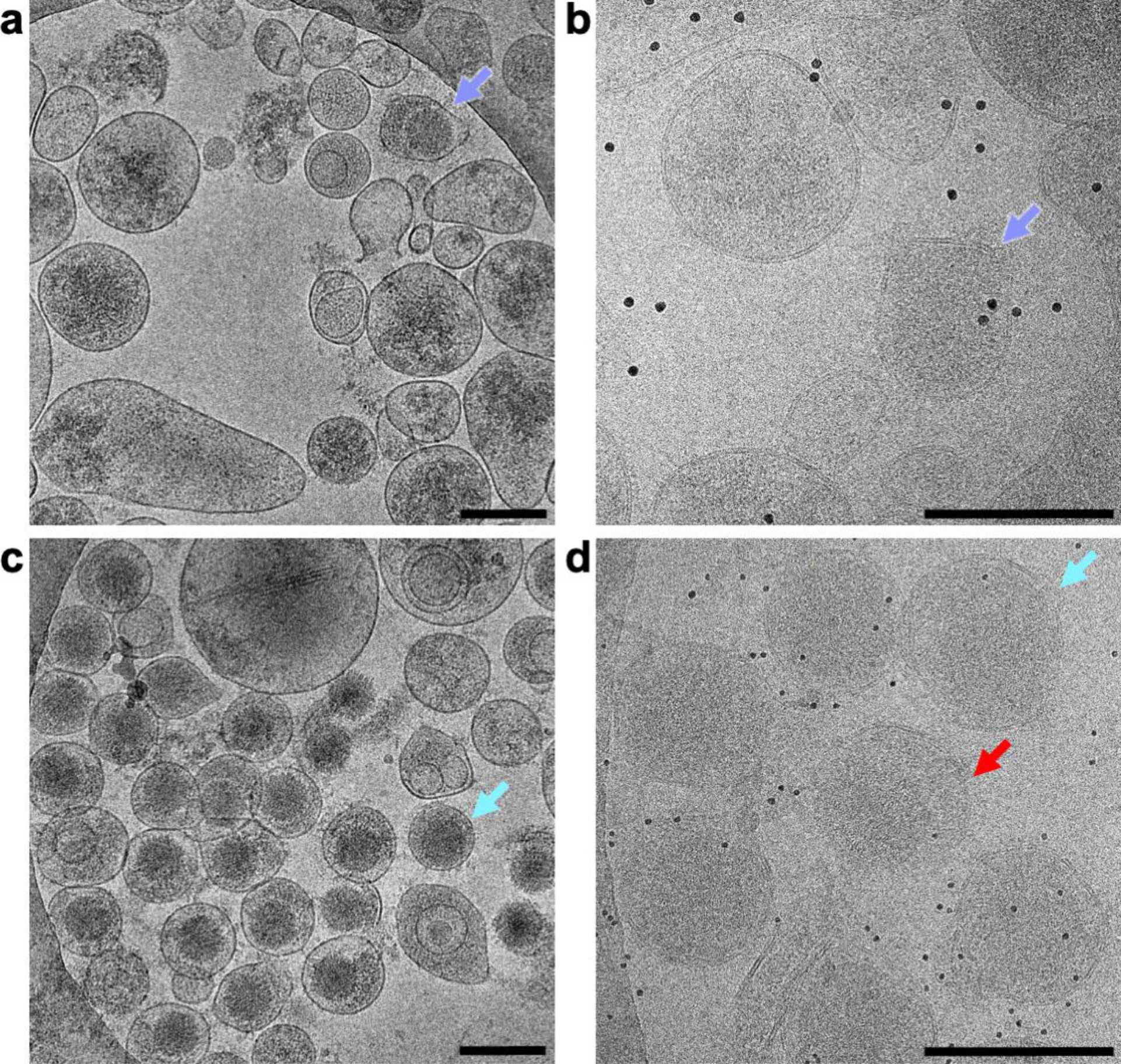
Isolated EBV and KSHV virion samples. (a) CryoEM screening image of isolated EBV virion sample. Intact EBV virion is indicated by arrow (orchid). Scale bar, 200 nm. (b) Low-dose image of EBV virion sample with 10 nm gold fiducials at a tilt angle of zero degrees for tomography. Intact EBV virion is indicated by arrow (orchid). Scale bar, 200 nm. (c) CryoEM screening image of isolated KSHV virion sample. Example of intact KSHV virions is indicated by arrow (sky). Scale bar, 200 nm. (d) Low-dose image of KSHV virion sample with 5 nm gold fiducials at a tilt angle of zero degrees for tomography. Example of intact KSHV virions is indicated by arrow (sky). Red arrow indicates broken virion. Scale bar, 200nm.

In addition to EBV, we isolated and imaged KSHV virions derived from the iSLK cell system (44). In contrast to the EBV sample, the KSHV sample has a much greater abundance of virions (Fig. 1c). Although vesicles are also present, they are less obstructive. In tilt series images of KSHV, we can also resolve individual bilayer leaflets and dsDNA contours (Fig. 1d). Because of the greater availability of virions in the KSHV sample relative to the EBV sample, we could collect a more robust dataset for KSHV virions.

We used cryoET to reconstruct the 3D structures of individual EBV and KSHV virions. Initial tomograms were reconstructed through conventional weighted back-projection with a simultaneous iterative reconstructive technique (SIRT)-like filter (Fig. 2a,e). In these conventional tomograms, we observe the capsid, tegument, and envelope of EBV and KSHV. The trace of the dsDNA genome within the capsid can be observed in some virions (Fig. S1). We are able to resolve the leaflets of the bilayer envelope, but the envelope appears distorted along the Z direction due to the missing-wedge problem inherent to cryoET (47). Features on the envelope and within the tegument layers are also difficult to discern due to poor contrast and high noise. For repeating particles of like features, subtomogram averaging is employed to overcome the missing-wedge problem while simultaneously increasing the signal-to-noise ratio and resolution of the resulting map (48). However, herpesvirus tegument and envelopes are pleiomorphic, so subtomogram averaging cannot be used to remedy missing-wedge artifacts and low contrast.

**Figure 2.**
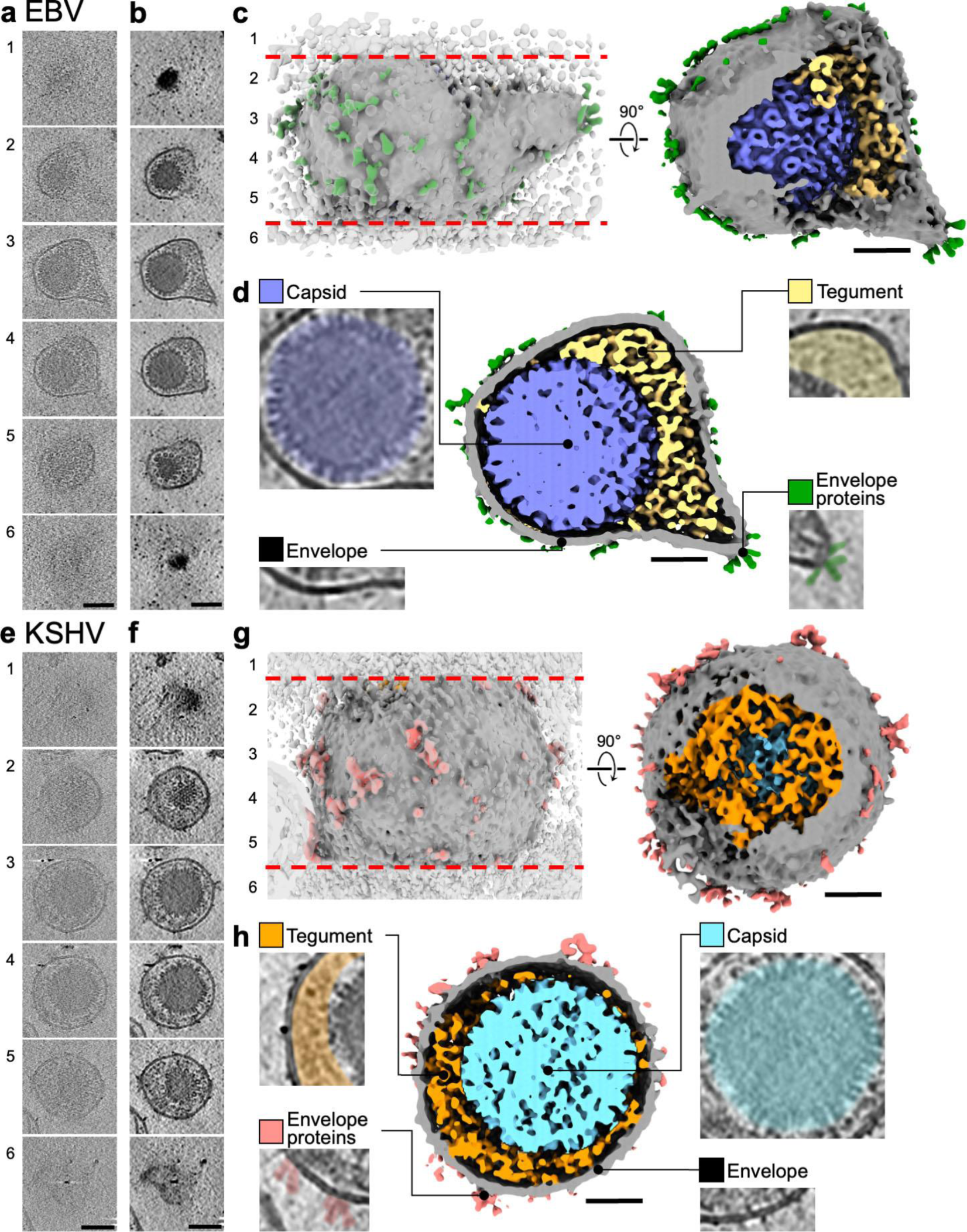
CryoET of EBV and KSHV. (a,e) Series of tomogram slices from weighted-back projection reconstruction with SIRT-like filter along z-axis of virion. Numbers correspond to location in surface rendering depiction (c,g). Scale bar, 80 nm. (b,f) Series of tomogram slices from isotropic reconstruction along z-axis of virion. EM density at top and bottom represent envelope, which indicates intact virion. Scale bar, 80 nm. (c,g) Surface renderings of virion reconstructions were cropped at the top and bottom poles (red lines) prior to segmentation to improve interpretability. Numbers indicate locations of corresponding tomogram slices (b,f). Scale bar, 40nm. (d,h) Sectional view of 3D surface rendering of virion colored by different components. Tomogram slices of corresponding virion component densities are indicated with their respective color. Scale bar, 40 nm.

To overcome these problems and to improve interpretability of our herpesvirus tomograms, we applied IsoNet, a deep learning-based software for the correction of missing-wedge artifacts and denoising (41) (Fig. S2). Representative IsoNet-processed tomographic slices of an EBV and KSHV virion show intact viral envelopes (Fig. 2b,f). In spite of the remaining noise in the map, we can identify a mostly smooth surface corresponding to the viral envelope (Fig. 2c,g). Small protrusions connected to the smooth envelope exterior are considered to be envelope proteins. In a clipped view of EBV and KSHV segmentation (Fig. 2d,h), the layered architecture of the nucleocapsid, tegument, envelope, and envelope proteins are demarcated. Individual capsomeres and envelope proteins are distinguishable in the tomographic slices and the surface rendering. Noticeably, the bulk of the tegument is offset to one side of the virion, as previously observed in HSV-1 and HCMV (36, 39). Because of pixel binning required by IsoNet, the gap between adjacent dsDNA can no longer be resolved, so the contour of the genome cannot be traced in those reconstructions (Fig. S2) (22). Although the KSHV and EBV reconstructions are of insufficient resolution for atomic model building, they nevertheless enable comparison between the two species of human gammaherpesvirus.

### Subtomogram averaging of nucleocapsids reveals distinct capsid-associated features in KSHV and EBV

To investigate the structure of the nucleocapsids, we performed subtomogram averaging of the EBV and KSHV icosahedral capsids (Fig. 3a,b). The EBV and KSHV capsids were reconstructed with icosahedral symmetry to ∼24 Å and 17 Å resolution, respectively (Fig. S3, S4). At these resolutions, the hexons, pentons, and triplexes are observed. Both of our EBV and KSHV icosahedral capsid reconstructions are similar to their respective cryoEM maps and can be fitted by the associated atomic model (29, 30, 38). Capsid vertex-specific component (CVSC) densities stemming from the triplexes adjacent to the pentons are observed in both the EBV and KSHV capsids (49–52), and their handed arrangement around the vertex validates the handedness of our subtomogram averaging reconstructions.

**Figure 3.**
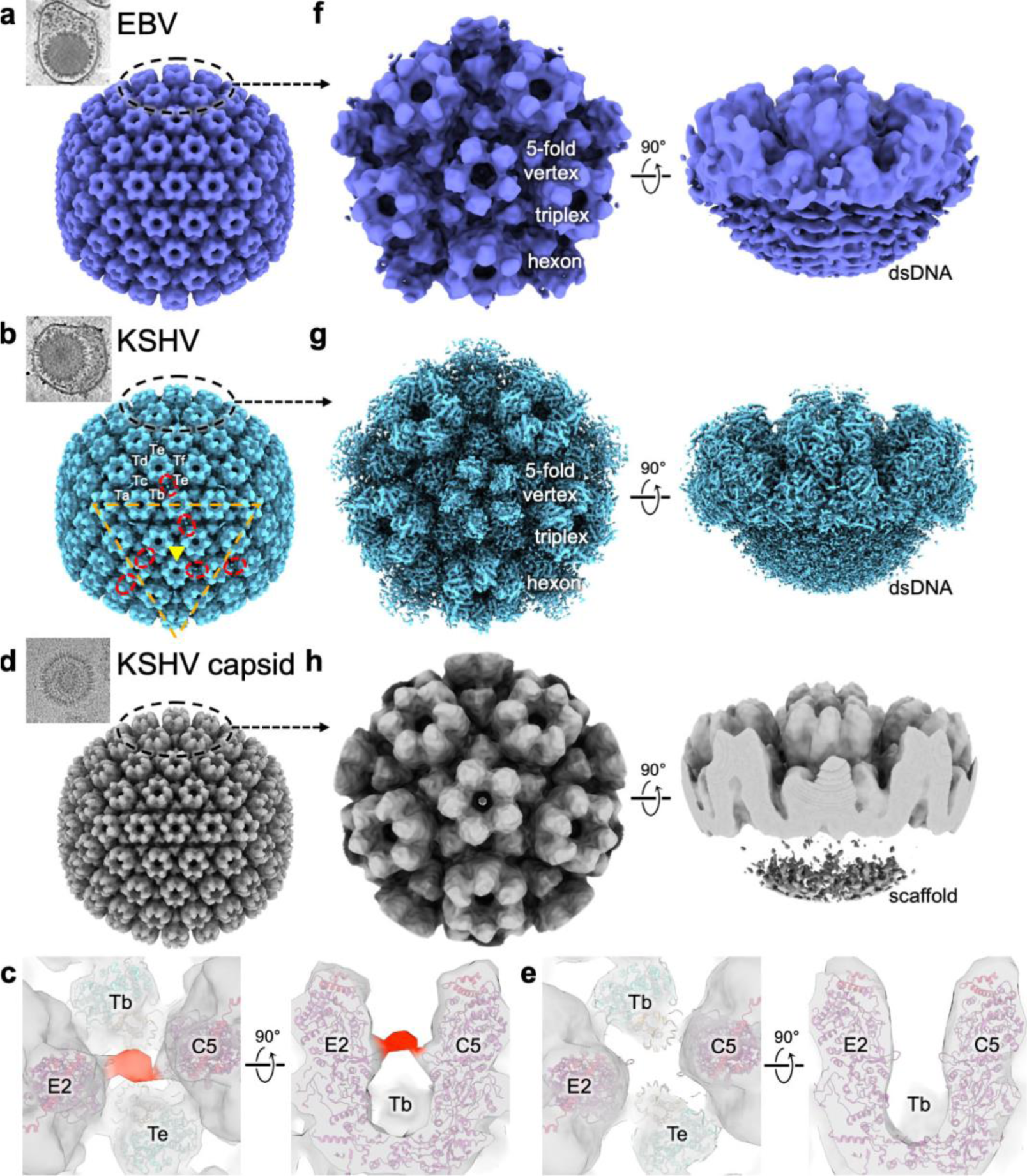
Subtomogram average reconstructions of EBV and KSHV capsid. (a) Subtomogram average of EBV capsid with icosahedral symmetry. Top-left shows tomogram slice of example EBV capsid from intact virion used for reconstruction. (b) Subtomogram average of KSHV capsid with icosahedral symmetry. Top-left shows tomogram slice of example KSHV capsid from intact virion used for reconstruction. Additional densities of unknown identity are indicated (red circle). Two-fold symmetric interfaces and three-fold symmetric axis are represented by orange lines and yellow triangle, respectively. Triplex nomenclature is annotated. (c) Overhead and side view of additional KSHV density from (b) indicated in red. Atomic models of neighboring triplexes (Tb, Te) and MCP/SCP (E2, C5) (PDB ID: 6B43) are fitted around the additional density. (d) CryoEM map of naked KSHV capsid devoid of tegument lowpass filtered to the same resolution as KSHV subtomogram average. Top-left shows cryoEM image of example naked KSHV capsid devoid of tegument used for reconstruction. (e) Overhead and side view of region shown in (c) for naked KSHV capsid. (f) Tomogram-guided sub-particle reconstruction of EBV vertex with C_5_ symmetry. (g) Tomogram-guided sub-particle reconstruction of KSHV vertex with C_5_ symmetry.

Viewing the KSHV capsid reconstruction at low contours, we observe additional densities spanning hexons at the 2-fold interface to hexons at the adjacent 3-fold axes between triplexes Tb and Te (Fig. 3b). These densities form a network that bridges the capsid facets. When models of the KSHV asymmetric unit containing MCP, SCP, and triplex proteins are fitted into the map, this density is unaccounted for and presumably belongs to some capsid-associated tegument protein that contacts loops of the adjacent MCPs at residues ^548^PGAR^551^ (Fig. 3c). This additional density on the KSHV capsid has previously been observed and was attributed to low-affinity secondary binding sites of CVSC protein ORF32 (53). The density is also present in other KSHV capsid cryoEM maps from intact virions (26, 38). However, it is absent in our cryoEM map of the naked KSHV capsid devoid of tegument (Fig. 3d,e). Additionally, this density is absent in our EBV capsid reconstruction (Fig. 3a) and previous cryoEM capsid reconstructions for EBV, HSV-1, and HCMV (27, 29, 30, 33, 35, 36), which possess ORF32 homologs, thus suggesting that ORF32 does not meaningfully bind the capsid between Tb and Te. Combined with recent findings on herpesvirus CVSC structure and assembly (26, 36), this density between Tb and Te likely does not belong to ORF32, and the component is specific to KSHV tegument.

To analyze the 5-fold symmetric capsid vertex, the localized reconstruction method for cryoEM was adapted to cryoET (27, 54). The icosahedral capsid reconstruction was used to calculate the locations of the capsid vertices, which were then locally extracted as discrete subvolumes for subtomogram averaging. Through this method, we determined localized reconstructions of the EBV and KSHV capsid 5-fold symmetric vertex at ∼21 Å and 7.1 Å resolution, respectively (Fig. 3f,g; Fig. S3,S4). Prior published atomic models of the EBV and KSHV asymmetric units fit their respective cryoET capsid vertex reconstruction (29, 30, 38), and distinct strands of the underlying dsDNA genome are observed. Additional densities around the capsid are visible in both the EBV and KSHV virion vertex but not in the naked KSHV capsid vertex (Fig. 3h), which indicates that these densities belong to the tegument.

Unlike empty capsids whose portal complexes are apparent (20, 33), the portal vertex of nucleocapsids cannot be directly observed due to obfuscation by the enclosed genome. To determine the location of the portal in the nucleocapsids, we performed classification on the capsid vertex reconstructions to separate the portal vertices from the penton vertices. Although we were unable to distinguish the EBV portal vertex from EBV penton vertex using this approach, 3D classification successfully distinguished the KSHV portal vertex (9.8 Å resolution) from the KSHV penton vertex (7.0 Å resolution) (Fig. 4a,b; Fig. S4). The KSHV portal vertex reconstruction contains 554 out of the 7,177 total extracted capsid vertices (7.7%), which is in line with the expected one portal vertex out of 12 capsid vertices (8.3%). In a clipped side view of the penton vertex reconstruction, we observe clear densities for the penton MCPs and underlying dsDNA genome (Fig. 4c). We cannot observe the penton MCPs in the KSHV portal vertex, and we instead see densities resembling the pORF19 portal cap decamer and pORF43 portal complex dodecamer (Fig. 4d) (26, 55). The clip turret between the portal cap and portal complex is not well defined, likely due to its plasticity (26).

**Figure 4.**
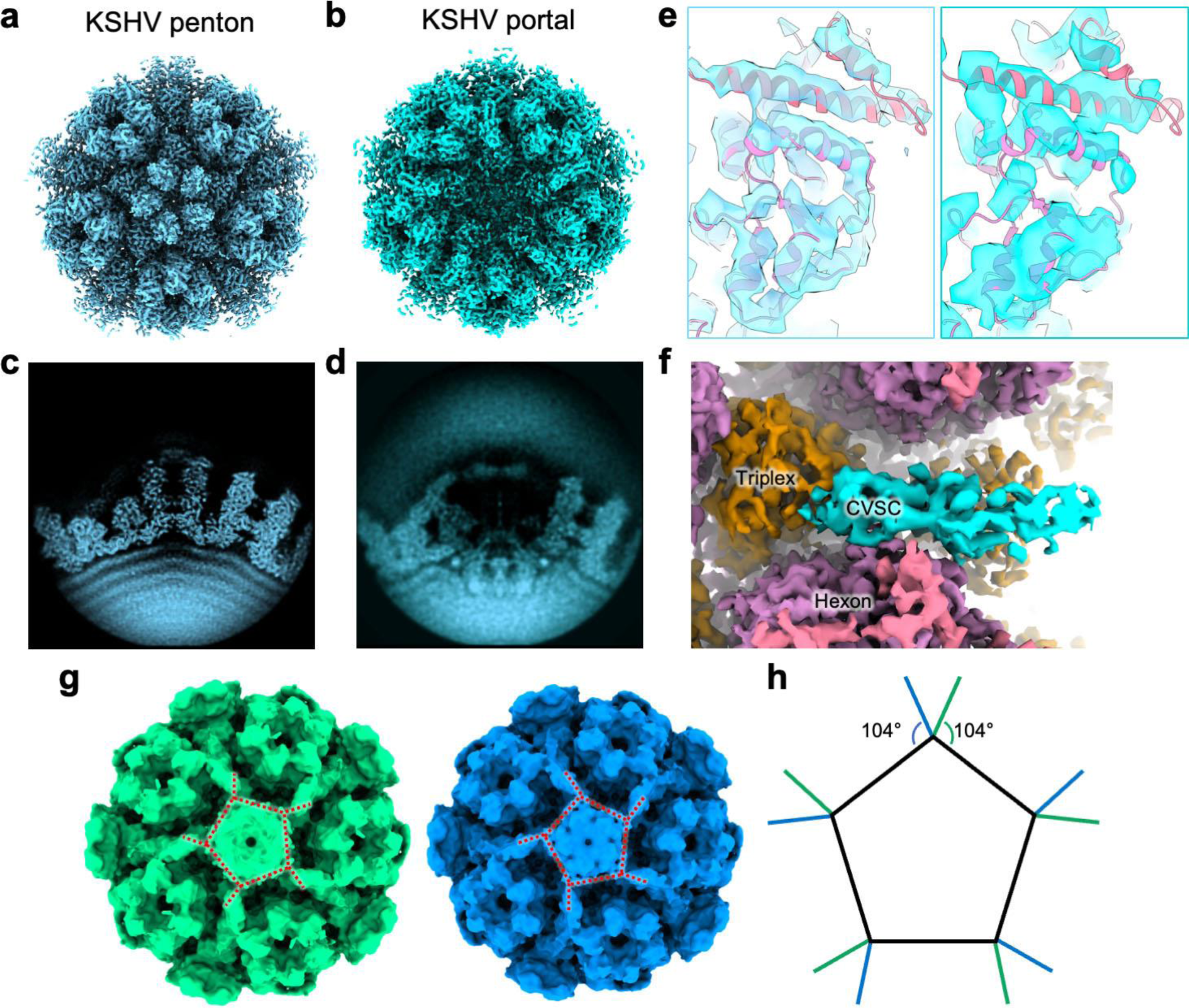
Tomogram-guided sub-particle reconstruction of KSHV penton and portal vertices. (a) Tomogram-guided sub-particle reconstruction of KSHV penton vertex with C_5_ symmetry (light blue). (b) Tomogram-guided sub-particle reconstruction of KSHV portal vertex with C_5_ symmetry (turquoise). (c) Side projection of KSHV penton vertex. (d) Side projection of KSHV portal vertex. (e) Secondary structure fit of KSHV MCP and SCP atomic model (PDB ID: 6B43) into cryoEM map of KSHV penton vertex (left) and KSHV portal vertex (right). (f) Segmentation of CVSC density (teal) in KSHV portal vertex. Triplex, MCP, and SCP are colored orange, purple, and pink, respectively. (g) Two classes of KSHV portal vertex (green, blue). Portal cap and binding site of CVSC are indicated by red dotted lines. (h) Overlay of portal cap and CVSC binding site schematic from (g). CVSC binding sites are colored by color of respective class (green, blue). The angles of CVSC and portal cap binding are indicated.

Although the penton vertex cryoEM map is of higher resolution than the portal vertex map and sufficient for fitting secondary structure features (Fig. 4e), the density for the CVSC in the penton map is poor compared to that in portal map (Fig. 4a,b,f). This difference in CVSC density suggests that the occupancy of CVSC at penton vertices is lower than at the portal vertex. Indeed, a prior cryoEM reconstruction without enforced symmetry shows that CVSC occupancy at vertices is directional with respect to the portal (26). Further classification of the portal vertex reveals heterogeneity of the pORF19 portal cap (Fig. 4g). Between the two classes, the binding of CVSC with the portal cap inverts handedness (Fig. 4g,h). In mature HCMV virions, variable configurations of portal cap association have been observed, including an inversion of the portal cap (36). Similar variability in portal cap orientation in KSHV suggests a conserved assembly pathway.

### Polarity and structures of the tegument layer

Due to the pleiomorphic nature of the tegument layer, tegument features cannot be averaged, so observations must be made from individual tomograms. We sorted tegument features into three previously described categories: inner tegument, outer tegument, and membrane-proximal tegument (56). To provide further context for tegument observations, we placed back the EBV and KSHV capsid reconstructions into the tomograms (Fig. 5a). From the positional information obtained from localized reconstruction of the capsid vertices, we also identify the locations of the capsid vertices in the tomograms. Since we are able to classify the portal vertices from penton vertices in KSHV, we also make use of the portal vertex to establish the orientation of the capsid within KSHV virions. Thus, capsid-relative tegument observations can be interpreted in relation to the capsid vertices for EBV and both the penton vertices and portal vertex for KSHV.

**Figure 5.**
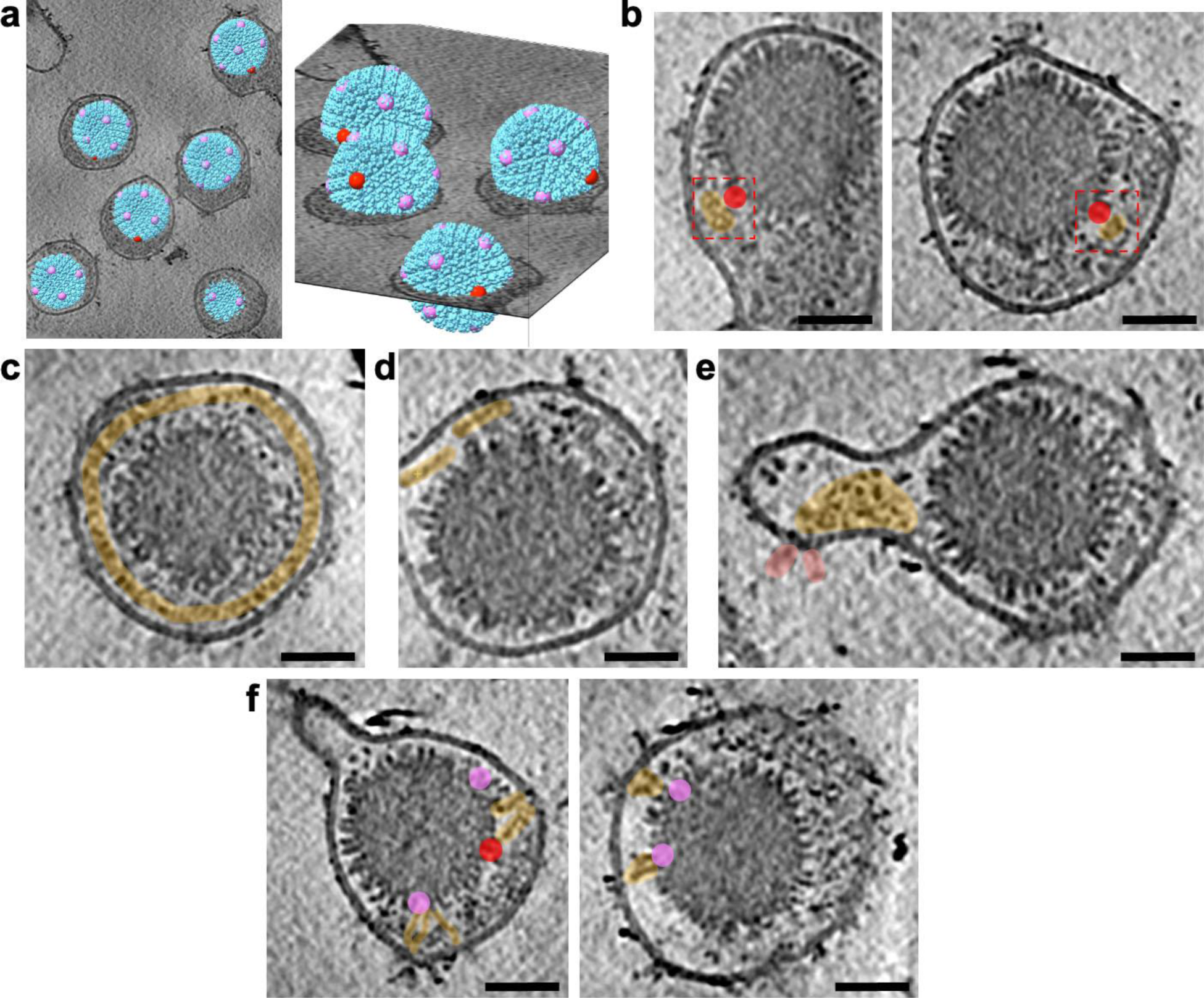
KSHV tegument in context of the portal and envelope. (a) Place back of the KSHV capsid 3D reconstruction (sky) into a tomogram. Portal vertex (red) and penton vertices (pink) for each virus are indicated. (b) Granular densities in tegument (orange) located near the portal (red). (c) Circumnavigation of envelope by discontinuous density resembling a dotted line (orange). (d) Long, thin densities in outer tegument bordering envelope. (e) Outer tegument (orange) associated with envelope protein (red) endodomains at a protrusion with few tegument densities elsewhere. (f) Long, thin densities spanning from capsid to envelope. Portal vertex (red) and adjacent penton vertices (pink) are indicated. Spanning densities are present in both presence (left) and absence (right) of tegument bulk. Scale bars, 40 nm.

For the majority of EBV and KSHV virions, the tegument is clustered to one side of the capsid, causing the capsid to be eccentrically located within the envelope (Fig. S5). This capsid eccentricity has previously been observed in HSV-1 and HCMV (36, 39) and has been recapitulated in our cryoET of HSV-1 and HCMV (Fig. S5a). For KSHV, whose portal vertices we could determine, we calculated a “portal-capsid-tegument angle” (PCT angle), a previously used proxy for quantification of the angular distribution of tegument relative to the portal vertex (Fig. S5b) (36). For KSHV, across 265 virions, 149 virions (56.2%) have a bulk of tegument within 40° of its portal vertex. This differs from previously published measurements for HCMV, where 38 out of 49 virions (77.6%) have a PCT angle of less than 40° (36). Although asymmetric tegumentation is a conserved feature of herpesvirus assembly, direction towards the portal vertex appears to vary, possibly due to different binding affinities of species-specific tegument components.

Distinct globular densities of ∼20 nm by 15 nm by 10 nm are observed in the outer tegument near certain EBV and KSHV capsid vertices (Fig. 5b). More specifically for KSHV, these globular densities have an increased propensity at the portal vertex. The globular densities are offset from the center of the vertex, which suggests that these densities may belong to tegument protein species that bind to CVSC or other capsid-associated inner tegument proteins. In addition to contacts between the tegument and capsid, noticeable areas of low density are present between the tegument and capsid (Fig. 5b). These areas of low density indicate low tegument association or lack thereof with the capsid.

Along the virion envelope of both EBV and KSHV, a disjointed layer of membrane-proximal tegument circumnavigates the interior (Fig. 5c). From an orthogonal view, the densities present as long strands arranged from end to end discontinuously (Fig. 5d) This layer is composed of densities with a diameter of ∼6 nm and varying lengths up to ∼50 nm. The diameter and filamentous nature of these densities suggest that they may be actin, which is found near secondary envelopment sites and could feasibly be incorporated within the virion during secondary envelopment (57–59). Similar actin-like densities have also been observed in HSV-1 (39). These filamentous components within the tegument may function as structural reinforcement of the envelope and in bridging the nucleocapsid to the host cortical actin during cellular entry.

Tegument also tends to congregate beneath regions of the envelope that contain envelope proteins, indicating interactions between the tegument and envelope proteins. This is most noticeable at protrusions with overall low proteinaceous tegument densities except for the regions containing clusters of envelope proteins, which instead have large densities below them (Fig. 5e). The large densities below clusters of envelope proteins are too large to be solely attributed to the endodomains of the envelope proteins, so they are likely composed of tegument proteins associated with the endodomains of envelope proteins and comprise the membrane-proximal tegument and outer tegument layers.

We observe thin densities spanning between the capsid and the envelope (Fig. 5f). The densities are ∼4 nm width and 15-30 nm long and the capsid end primarily localizes near penton vertices. Long proteins stemming from the nucleocapsid have previously been observed in HSV-1 (pUL36) and HCMV (pUL48) (36, 60, 61). These proteins are homologous to BPLF1 and ORF64 in EBV and KSHV, respectively. Crosslinking MS experiments in HCMV have indicated tegument species that connect to both the capsid and the envelope (56), so similar spanning tegument proteins may be present in other herpesviruses, including EBV and KSHV. The major spanning protein in HCMV (UL32/pp150) does not have an equivalent in other herpesviruses, which suggests that EBV and KSHV may possess unique proteins that share a similar function as a scaffold for recruitment of the herpesvirus layers.

### Morphology of the envelope and volume of the virion

EBV has an average volume and surface area of 2.6×10^6^ nm^3^ and 1.2×10^4^ nm^2^, respectively, and KSHV has an average volume and surface area of 2.7×10^6^ nm^3^ and 1.1×10^4^ nm^2^, respectively (Fig. S5c,d). These average volumes and surfaces areas are similar to those of HSV-1 and HCMV. Both EBV and KSHV envelopes are generally round with deviations from sphericity caused by the asymmetric association of tegument around the capsid. Additionally, some virions have protrusions of varying prominence (Fig. 5e,f). Some protrusions are filled with tegument while others are devoid of proteinaceous density. Although we cannot exclude the possibility that protrusions result from the purification process, previous studies of herpesvirus virions within the cytoplasm of infected cells also possess them (43, 62–64), which suggests a physiological morphology. Virions isolated by ultracentrifugation remain infectious (Fig. S6), indicating that structural studies of herpesvirus virions prepared in this manner retain features that reflect infectivity.

Some regions of the envelope with spherical curvature do not have underlying membrane-proximal tegument (Fig. 5b,f). This typically occurs when the capsid is in direct contact with the envelope and occludes space for a tegument layer. Noticeably, the void of tegument also presents as a distinct gap between the envelope and the capsid as though the capsid has been pulled away from the envelope (Fig. 5f). The envelopes around these voids retain their curvatures, which suggests that these regions are still internally supported. Serendipitously, the lack of bulk tegument in these void regions permit clearer observation of the tegument proteins that remain, including the thin densities that span the capsid and the envelope (Fig. 5f). Envelope proteins appear on the opposing side of where these thin densities contact the envelope, which suggests that some species of envelope proteins function as anchors for the thin densities to bind. Due to their presence outside of the bulk tegument, we infer that these thin densities bind the capsid external of the other tegument species and possibly function in tethering the membrane to the capsid.

In addition to intact virions, we also observe EBV and KSHV particles that deviate from the conventional depiction of herpesvirus. These include enveloped capsids with invaginations and with much larger volume than average (Fig. S7). We also observe envelopes containing two capsids. These indicate either potential fusion events between virion and vesicle or between virion and virion. We observe a potential intermediate state of virion fusion with another virion as their envelopes are connected by a thin pore (Fig. S7), which suggests that virion-virion fusion may occur. Alternatively, an envelope containing two capsids may be the result of mispackaging of two capsids simultaneously during secondary envelopment (Supplemental Fig. S7). These deviations from the “norm” emphasize the irregularities in herpesvirus assembly.

### Herpesvirus envelope protein organization and correlation with infectivity

To characterize the envelope proteins of EBV and KSHV, we quantified them and noted their arrangements on the envelope (Fig. 6). Notably, both EBV and KSHV appear sparsely decorated with envelope proteins compared to HSV-1 and HCMV (Fig. 5a). Counting external densities of at least 5 nm protruding from the envelope surface, EBV and KSHV have two- to threefold fewer presenting envelope proteins than HSV-1 and HCMV (Fig. 5b). Because envelope proteins are integral for infection, a greater density of them on the envelope should increase the likelihood of encountering a host receptor and initiating membrane fusion for cell entry. Through a GFP-based infection assay, we determined that the number of presenting envelope proteins correlates with infectivity (Fig. 5c). KSHV has an eightfold greater infection rate compared to EBV in the infection assay despite both having similar counts of envelope proteins. This could be attributed to the greater binding affinity for EphA2 by the KSHV gH/gL complex compared to that of EBV (65). However, we cannot establish causation between envelope glycoprotein density and infectivity due to variables in the experimental design, including differences in timing of post-infection expression and host tropism.

**Figure 6.**
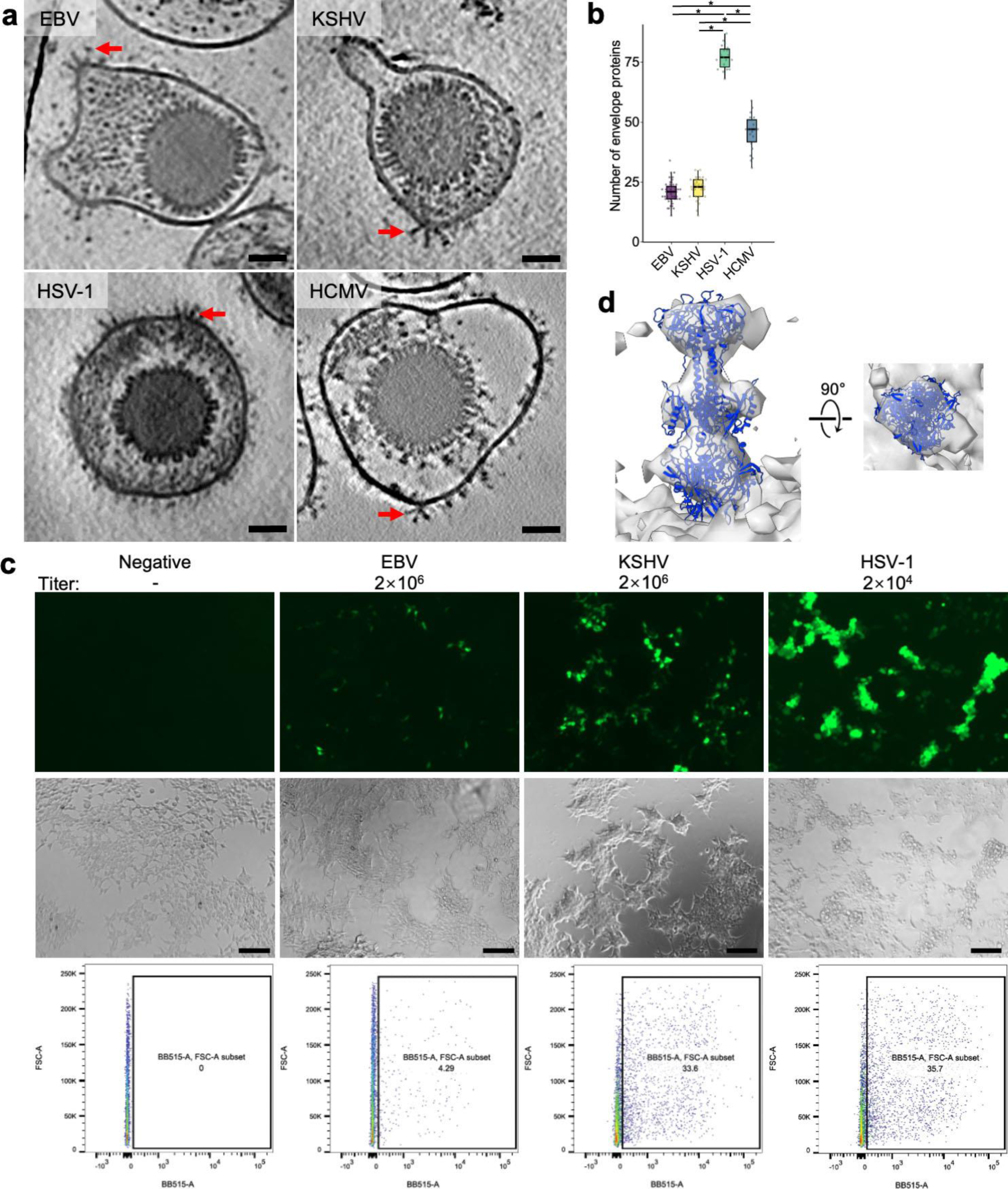
Envelope protein organization and correlation with infectivity. (a) Comparison of tomogram slices between EBV, KSHV, HSV-1, and HCMV. Clusters of envelope proteins are indicated by red arrow. Scale bar, 40 nm. (b) Count of envelope proteins per virion for indicated herpesvirus species. Statistical analysis used ordinary one-way analysis of variance (ANOVA) with Tukey’s test for multiple comparisons (*, P < 0.05). (c) Relative initial infectivity assay of HEK293T cells by EBV, KSHV, or HSV-1 with titer measured as genome copy number by qPCR. Top row: GFP fluorescence indicates infected cells. Middle row: Light microscopy image of view from fluorescence images. Bottom row: Count of infected cells by flow cytometry. Scale bar, 150 μm. (d) Sub-tomogram average reconstruction of large envelope proteins from clusters in KSHV. Postfusion gB atomic model (blue; PDB ID: 3FVC) fits the cryoET density.

We noticed that envelope proteins tended to cluster on the envelope. With the aid of deep learning-enhanced isotropic reconstructions, we better resolve the envelope protein clusters and identify their arrangements as triplets, sets of three envelope proteins (Fig. 5a). Subtomogram averaging of the envelope proteins in these triplet clusters identifies them as postfusion gB (Fig. 5d). Envelope glycoproteins tend to cluster on protrusions, but gB units along these protrusions tend to be in the postfusion conformation. This preference for the postfusion state can be attributed to the increased curvature at these regions inducing conformation change (66, 67). Alternatively, the prefusion state of gB may be present but less identifiable than the postfusion state due to its less distinct shape and the low contrast of the tomograms. Additionally, the gB members of triplets are angled relative to the envelope like HIV-1 gp160 spike and SARS-CoV-2 spike (68, 69). Like the other spike proteins, the angle of gB may facilitate exploration of its envelope to engage with its host receptor (69).

## Discussion

CryoET reconstructions of EBV and KSHV with spatially determined capsids enable contextualized observations of their pleiomorphic tegument and envelope. Both EBV and KSHV share similar overall morphology for the dsDNA genome and capsid, but we observe differences envelope protein distribution. The tegument is not fully amorphous as commonly depicted and instead contains distinct structural features (Fig. 5). Tegument-mediated contacts between the envelope and the capsid suggests an interlayer dynamic for the tegumentation and envelopment pathways during herpesvirus assembly. The envelope proteins are further organized into distinct clusters that could mediate function, and their relatively low prevalence on EBV and KSHV could explain their infectivity (Fig. 6).

Intracellular herpesvirus virions have been observed with deviations from sphericity, including protrusions (43, 70), which suggests that these perceived flaws may instead be a feature. Protrusions and other deviations from sphericity increase the surface area of an envelope for a given internal volume, which increases the potential interaction sites with a host cell. We notice that deviations from sphericity vary between herpesvirus species (Fig. S5a), which suggests that the pliability of the envelope and its composition may differ between them. The tegument may also be involved with maintaining the shape of the envelope by anchoring it to the capsid. This is most noticeable in HCMV (40), where the envelope protrudes when underlying tegument is lacking (Fig. 5). The outer tegument may not be tightly associated with the capsid in order to disperse inside the cell and prepare the host environment for viral expression and assembly.

In prior cryoET studies of HSV-1 infection, envelope proteins are not observed to be recruited to the entry site, suggesting that few gB units are necessary to catalyze entry (71). In our tomograms, those few gB units appear to be clustered on the envelope into functional units to facilitate entry (Fig. 5a). The clustering of gB may aid their membrane fusion function, which has been observed in other fusogenic proteins. SNARE proteins require three contact points to form a pore (72). The homologous vesicular stomatitis virus G protein requires a certain number of copies clustered together to trigger hemifusion (73). The assembly of these gB clusters remains to be determined but may occur through directed localization by trans-Golgi network, lipid rafts, tegument, or cytodomain of gB (74–76).

The different herpesvirus species have different distributions of proteins displayed on the envelope (Fig. 5a,b), which may explain the differing neutralizing antibody profiles between herpesviruses (77–87). Gammaherpesvirus have low immunogenic response to gB as measured by anti-gB from virion immunization or patient derived sera (81, 88, 89). The herpesvirus tomograms from this study suggests that the low prevalence of gB on the surface of EBV and KSHV may be the cause of the weak gB response by the immune system. HSV-1 and HCMV may have higher anti-gB response from sera than their gammaherpesvirus relatives due to this greater gB presence on their envelopes. Although anti-gB antibodies have neutralizing activity against EBV and KSHV like they do against HSV-1 and HCMV (90, 91), their generation via attenuated EBV or KSHV virions or virus-like particles may not be as effective compared to similar processes of neutralizing antibody generation for HSV-1 or HCMV. As such, EBV or KSHV gB overexpressed and displayed on vesicles or nanoparticles may better elicit production of antibodies with broad neutralizing activity than attenuated virions.

The most recent 3D structural characterization of an intact gammaherpesvirus virion is a cryoET study of MHV-68 (43), a model system for EBV and KSHV due to its relative ease of culture (92). In MHV-68, a distinct boundary just above the capsomere towers demarcates the inner tegument layer and distinguishes it from the outer tegument layer (43). In our tomograms of EBV and KSHV, this boundary manifests instead as a lighter region of spaces between the capsomeres and a darker region of the bulk tegument (Fig. 5), blurring the distinction between inner and outer tegument. Occasionally, the tegument bulk infiltrates the space between capsomeres, suggesting an intermingling between the capsid-associated inner tegument and the outer tegument. This difference in visualization could be attributed to improved tomogram-level resolution due to hardware and software advancements since the previous study.

Although cryoET has enabled observation of structures and features of EBV and KSHV, identification of individual proteins is inhibited by the limited resolution of the tomograms. The resolution is primarily limited by poor signal-to-noise ratio from the need to work around radiation damage of the sample (93). This problem is further exacerbated by the inherent thickness of the sample to accommodate the large virions (94). The volta phase plate has been used to obtain higher contrast (40), but they contradictorily reduce the high-resolution signal (95). Deep learning enhancement currently down samples the tomograms and prevents recovery of information for small features. Future developments in cryoET technology are required to increase attainable resolution of thick samples at the tomogram level in order to truly understand herpesvirus architecture and underlying molecular sociology.

## Materials and Methods

### EBV culture

7.5×10⁵/mL of B95-8 cells (ATCC CRL-1612) infected with EBfaV-GFP (recombinant EBV reporter virus expressing GFP) (96) were grown in 100 mL RPMI 1640 (Corning 10-040-CV) with 10% FBS, penicillin-streptomycin, and 30 ng/mL 12-*O*-tetradecanoylphorbol-13-acetate (TPA; Sigma) at 37°C and 5% CO₂ for 4 days. The supernatant was collected by centrifugation at 1,500 RPM for 10 minutes.

### KSHV culture

KSHV virions were prepared as previously described with slight modification from iSLK-KSHV-BAC16 cells, received as a gift from Dr. Jae U. Jung (44, 53). In brief, iSLK-KSHV-BAC16 cells were cultured in Dulbecco’s Modified Eagle Medium (DMEM; Corning 10-017-CV) supplemented with 10% fetal bovine serum (FBS; 35-010-CV) and 100 U/mL of penicillin-streptomycin, 1mg/mL puromycin (Invivogen ant-pr-1), 250 μg/mL G418 (Invivogen ant-gn-1), and 1,200 mg/mL hygromycin B (Invivogen ant-hg-1). Cells were cultured to 80-90% confluency in 24 15 cm tissue culture dishes. Then, KSHV lytic replication and virion production were induced by treatment with 1 mM sodium butyrate and 5 μg/mL tetracycline for five days, after which 720 mL of media containing KSHV virions were collected.

### HCMV culture

HCMV virions were prepared as previously described with slight modification (35). Human fibroblast MRC-5 cells (ATCC CCL-171) were seeded on 18 T175 flasks in Eagle’s Minimum Essential Medium (EMEM; ATCC 30-2003) supplemented with 10% FBS (R&D Systems S11150H) and 100 U/mL of penicillin-streptomycin (Gibco 15140122) and cultured until they reached ∼90-100% confluency. The cells were then infected with HCMV (AD169 strain, ATCC VR-538). The growth media were replaced 3 days after infection, and the media containing secreted HCMV virions were collected at 7 days post-infection.

### HSV-1 culture

HSV-1 virions were prepared as previously described with slight modification (23). Vero cells (ATCC CCL-81) were seeded on 12 T175 flasks in DMEM (ATCC 30-2002) supplemented with 10% FBS and 100 U/mL of penicillin-streptomycin. At 100% confluence, the cells were infected with HSV-1 (KOS strain, ATCC VR-1493). The media containing secreted HSV-1 virions were collected 3 days post-infection.

### Herpesvirus virion isolation

Herpesvirus virions were isolated as previously described with slight modification (97). Infectious media was clarified by centrifugation at 10,000 × g for 20 min at 4 °C twice. Virions were then pelleted using an SW28 rotor (Beckman) at 80,000 × g for 1 h at 4 °C. The supernatant was discarded and 50 μl of cold phosphate-buffered saline (PBS) was added to the pellet in each tube. The pellets were allowed to soften for 16 h on ice in a 4°C cold room before gentle resuspension by tapping. About 0.7 ml of combined pellet suspension from all ultracentrifuge tubes were loaded on 15–50% linear gradient of sucrose made in PBS and spun in SW41 (Beckman) rotor at 80,000 × g for 1 h at 4 °C. About 1 ml of the visible virus-containing band was collected into a new tube and mixed with 10 ml of sterile PBS solution in order to dilute sucrose. EBV did not have a visible band and was instead fractionated in 1 mL aliquots based on the band locations of other herpesviruses. The virions were concentrated at 80,000 × g for 1 h at 4 °C and 20 to 40 μl of PBS was added depending on the pellet size. After softening and gentle resuspension, the isolated virions were recovered and immediately used for cryoET sample preparation without freezing.

### CryoET sample preparation and image acquisition

Virion samples were screened on an FEI Tecnai TF20 equipped with a 4k × 4k TVIPS F415MP CCD detector. To prepare EM grids for cryoET imaging, 3 μl of diluted virions were mixed with 5- or 10-nm gold fiducials and applied to Quantifoil R2/1 Cu 200 grids and plunge-frozen in liquid ethane-propane mixture using a Vitrobot mark IV (FEI Thermo Fisher Scientific) set to 100% humidity, 4 °C, and blot time of 5 s (98).

Imaging was performed with a Titan Krios G1 (FEI Thermo Fisher Scientific) operated at 300 kV, and movies were recorded using a Gatan K3 detector with an energy filter slit width of 20 eV. Tilt series were collected from −60° to 60° in 3° increments using a dose-symmetric scheme (99) using SerialEM (100) at 42,000 × magnification (1.044 Å/pixel super-resolution) and a target defocus of −5 μm or at 64,000 × magnification (0.690 Å/pixel super-resolution) and a target focus of −4 μm with a total of dose of 120 e−/Å^2^. Individual tilts were recorded with constant beam intensity in 10 frames. An HCMV dataset was collected with a Volta phase plate conditioned to induce an initial phase shift of π/3 and a defocus target of −0.6 μm. Data collection statistics are listed in Table S1.

### Tomogram reconstruction and subtomogram averaging

Each movie stack was drift-corrected using MotionCor2 (101) and defocus values were determined with CTFFIND4 (102). Tilt series alignment via gold fiducials and tomogram reconstructions were performed in IMOD with a binning factor of 4 (103). IsoNet isotropic reconstructions with a binning factor of 8 were used for particle picking and observations of non-repeating features (41). Capsid reconstructions of EBV and KSHV were performed in RELION-4 with a binning factor of 5 and 6, respectively (45). Tomogram-guided sub-particle reconstruction of the EBV and KSHV vertices at binning factor 2 was done as previously described for single particle cryoEM with slight modification for tomography data (27, 54). The formula for defocus change was instead used to shift the Cartesian position z. RELION-4 accounts for the depth-of-focus problem by using a defocus gradient for tomogram CTF correction (45). Classification of the EBV and KSHV vertices was performed with a cylindrical local mask focused over the 5-fold symmetry axis. For KSHV, one class with 7.7% of the particles contained portal complex features. Individual portal particles were placed back in their respective tomograms to verify that each parent capsid contained at most one portal. Redundant particles were removed based on _rlnMaxValueProbDistribution score. No distinct portal class was identified for the EBV particles. Subtomogram averaging of clustered envelope proteins was performed in PEET with the tilt orientation of particles coarsely determined using stalkInit (104). Visualization, segmentation, and measurements were performed in IMOD, ChimeraX, and ArtiaX (103, 105, 106).

### Infection assay

Infectious media was filtered using a 0.45 μM membrane (Millipore SE1M003M00), aliquoted into 1 mL portions, and stored at −80 °C for infection experiments. For testing infectious viability after ultracentrifugation, aliquots were centrifuged at 80,000 × g for 30 minutes at 4°C, and the pellets were resuspended in 100 μL of 10% FBS DMEM prior to infection.

Prior to infection, the viral titer was determined by qPCR. The DNA of EBV, KSHV, and HSV-1 was extracted using QIAamp DNA Kits (Qiagen) following the manufacturer’s instructions. The DNA copies of each virus were quantified using qPCR. This was based on the standard curve of the gB gene for each virus, utilizing specific gB plasmids for EBV, KSHV, and HSV-1. The original gB gene copies were calculated by dividing the plasmid weight by its molecular weight and then multiplying by the molar constant. The plasmids were serially diluted two-fold. Both virus DNA and diluted plasmids were subjected to qPCR. A linear standard curve was generated by plotting gB copies in plasmid against the Ct values from qPCR. Virus DNA copies were then calculated from the standard curve using the Ct values of the viral DNA. The following primers were used for real-time PCR: EBV, Forward: 5’- CAGCCAGACGGAGCTCTATG-3’, Reverse: 5’-AGAAGTCGAAGGGGCTGTTG-3’; KSHV: Forward: 5’-CTCGAATCCAACGGATTTGAC-3’, Reverse: 5’-TGCTGCAGAATAGCGTGCC-3’; HSV-1: Forward: 5’-CCGACCTCAAGTACAACCCC-3’, Reverse: 5’- GTAGCCGTAAAACGGGGACA-3’.

HEK293 cells (7.5×10^4^ cells per well) were seeded in 100 µL of 10% FBS DMEM medium in a 48-well plate and infected on the second day with the indicated amounts of EBV, KSHV, and HSV viruses in 100 µL of 10% FBS DMEM. Forty-eight hours post-infection, the cells were either visualized and captured using an EVOS fluorescence microscope or analyzed by flow cytometry.

### KSHV capsid isolation and cryoEM

KSHV capsid isolation, data collection, and data processing was performed as previously described (21).

## Data availability

CryoET tomograms have been deposited in the Electron Microscopy Data Bank under accession numbers EMD-XXXXX. The data that support this study are available from the corresponding author upon reasonable request.

## Acknowledgements

We thank Ana L. Alvarez-Cabrera and Zhu Si for assistance with HSV-1 sample preparation and imaging, Yun-Tao Liu for assistance with IsoNet, and Jonathan Jih and Alexander Stevens for insightful discussion. This research was supported in part by grants from the National Institutes of Health (R01DE025567 to Z.H.Z and T.-T.W., R21AI175798 to R.L., R01DE027901 to Y.Y.), grant IRG-15-173-21 (J.C.) from the American Cancer Society, a supplement from P30CA060553 (R.L.), and the Third Coast Center for AIDS Research pilot award (J.C.). J.Z. acknowledges support from an Interdisciplinary Training in Virology and Gene Therapy training grant (NIH 5T32AI060567). We acknowledge the use of resources in the Electron Imaging Center for Nanomachines supported by UCLA and grants from NIH (S10RR23057, S10OD018111, and U24GM116792) and NSF (DBI-1338135). We also acknowledge use of resources from the UCLA AIDS Institute, the James B. Pendleton Charitable Trust, and the McCarthy Family Foundation.

## Contributions

Z.H.Z., R.L., T.-T.W., R.S., and Y.Y. initiated and supervised the research. J.Z. prepared samples, recorded cryoEM and cryoET data, processed and interpreted the data, and prepared the manuscript. J.C. cultured EBV samples and performed infection assays. H.H. cultured KSHV samples. S. Liu prepared HSV-1 samples and obtained the HSV-1 tomograms. S. Liao processed KSHV data. All authors reviewed and edited the manuscript.

---

**Supplementary Figure S1.**
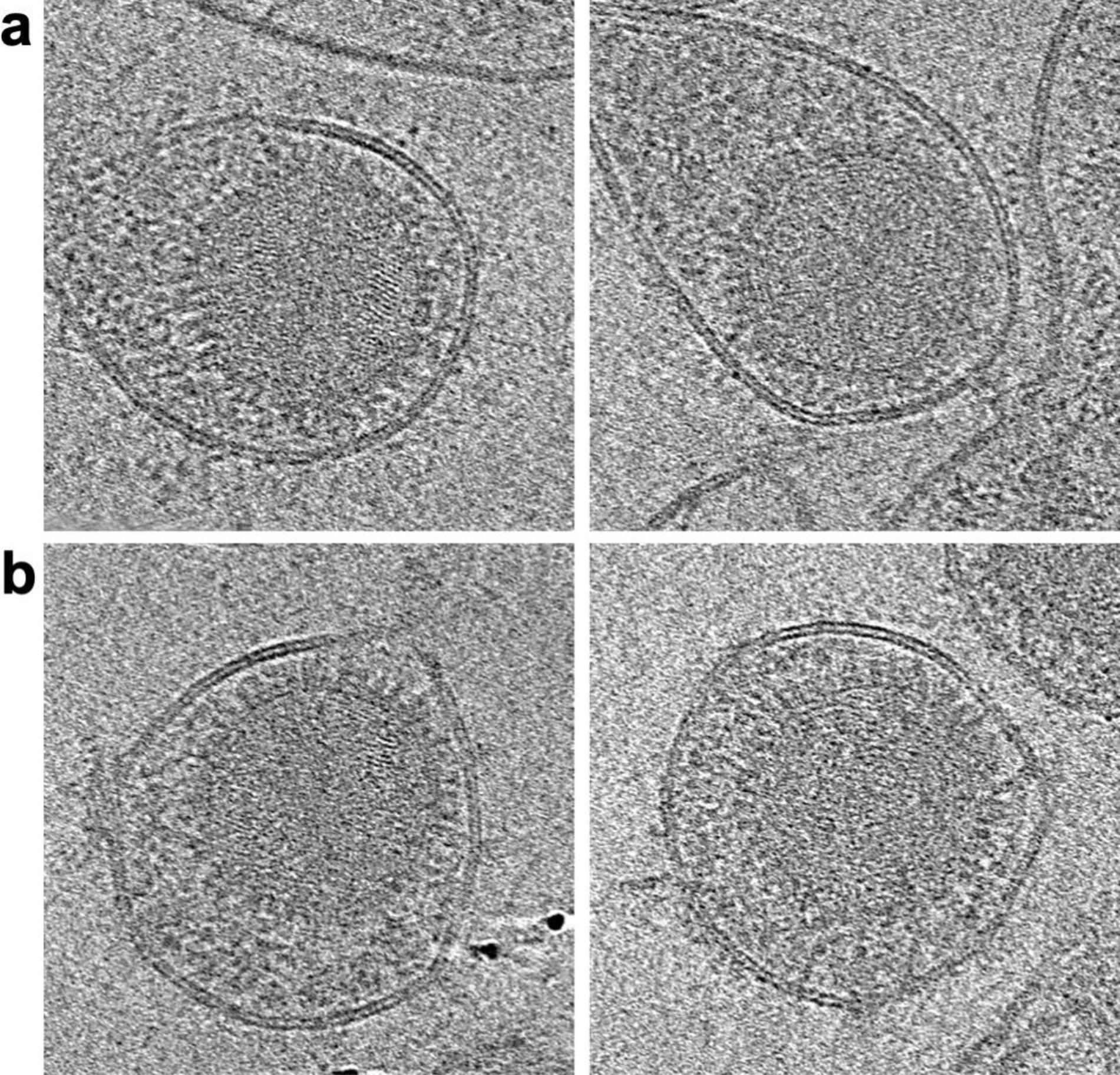
Tomograms capture genome strand contours. (a) Weighted back-projection with a simultaneous iterative reconstructive technique (SIRT)-like filter tomogram slice of EBV virion with genome oriented latitudinally (left) and circumferentially (right). (b) SIRT-like filter tomogram slice of KSHV virion with genome oriented latitudinally (left) and circumferentially (right).

**Supplementary Figure S2.**
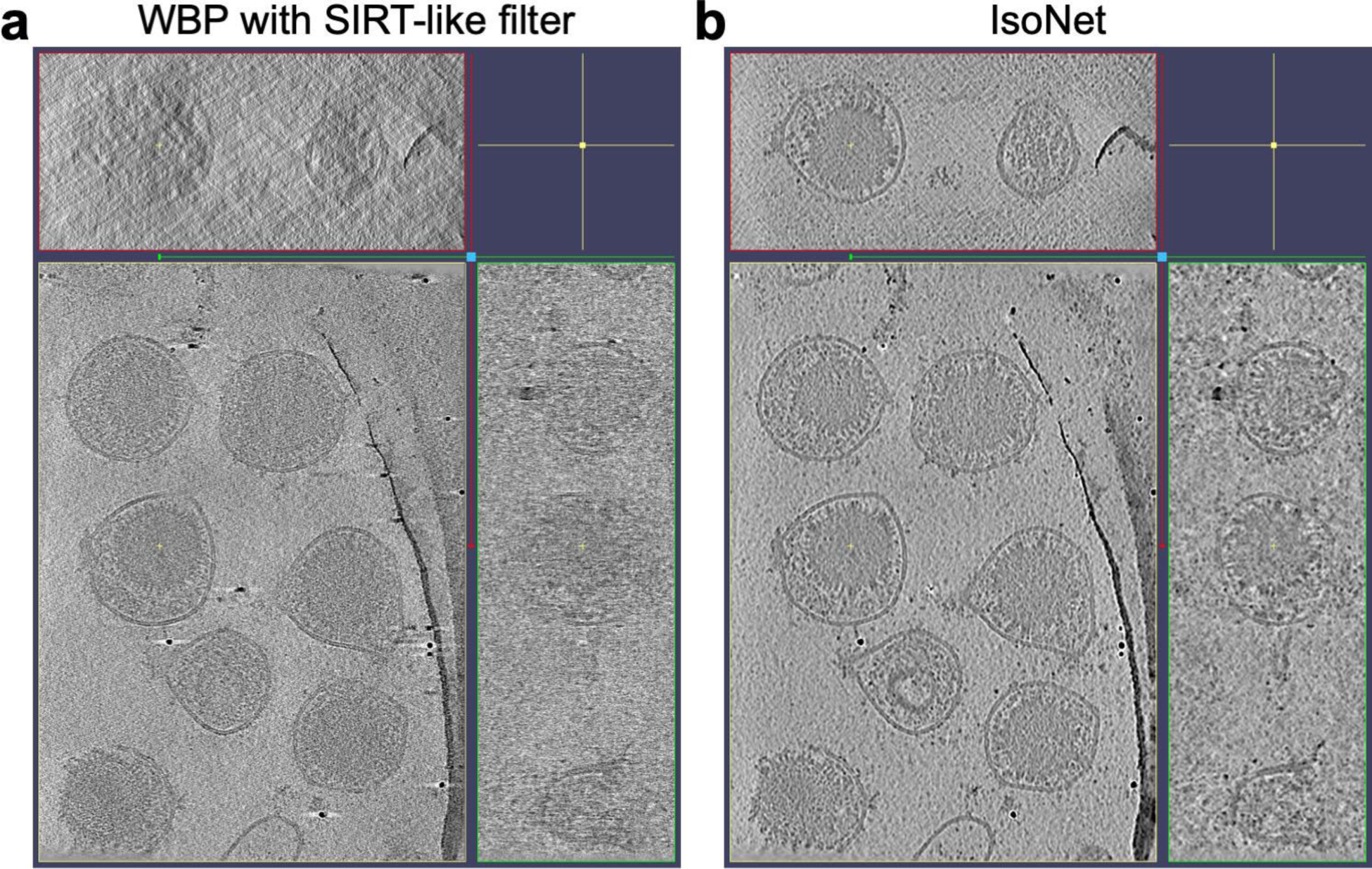
IsoNet processing of KSHV tomogram. (a) Weighted back-projection with a simultaneous iterative reconstructive technique (SIRT)-like filter tomogram displayed in XY, XZ, and YZ planes. (b) IsoNet processed tomogram displayed in XY, XZ, and YZ planes.

**Supplementary Figure S3.**
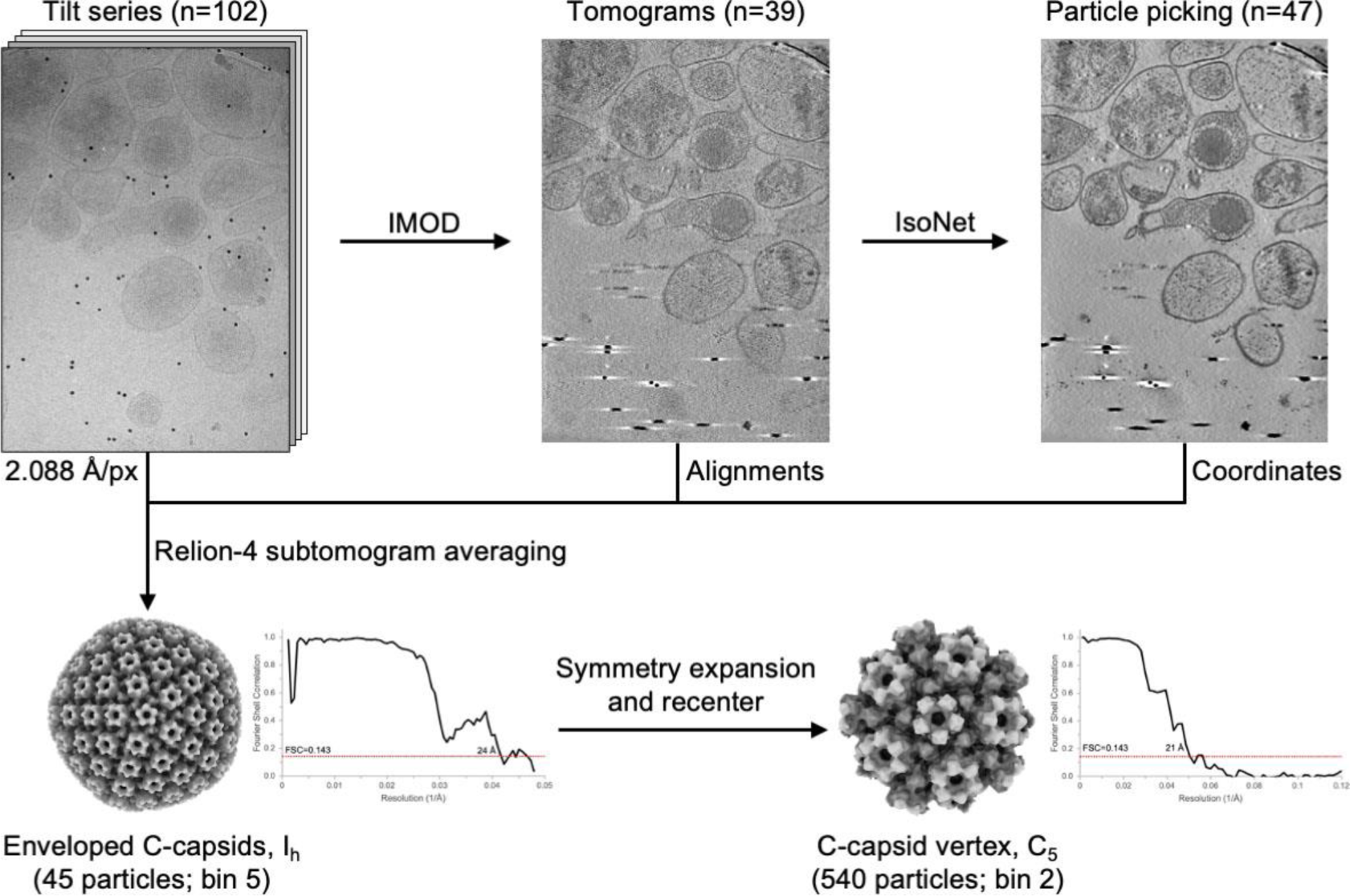
EBV processing workflow. Flowchart shows processing pipeline of tomogram-guided sub-particle reconstruction and reconstruction to resolve capsid vertex from icosahedral capsid.

**Supplementary Figure S4.**
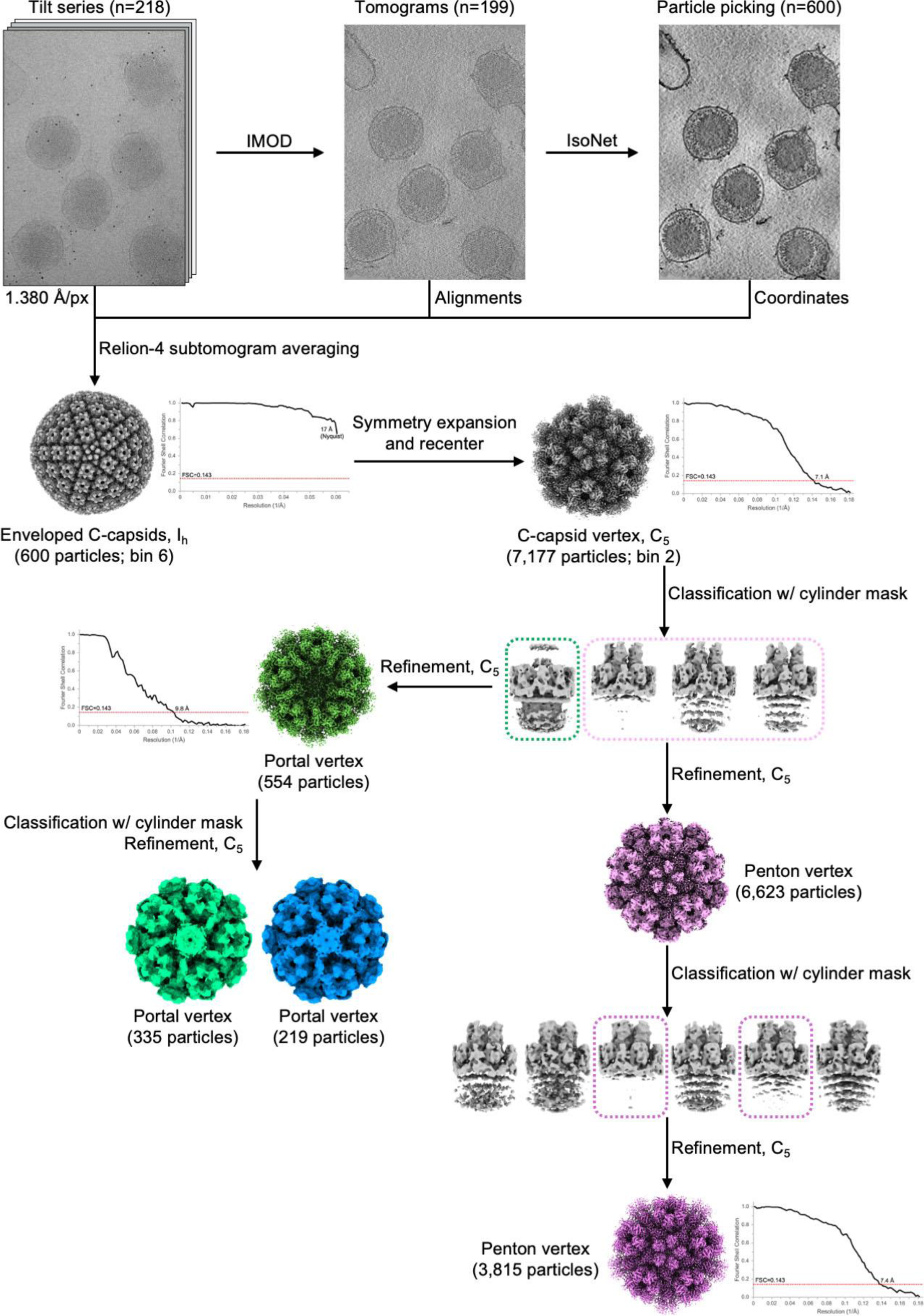
KSHV processing workflow. Flowchart shows processing pipeline of tomogram-guided sub-particle reconstruction and reconstruction to resolve capsid vertex from icosahedral capsid. Subsequent focused classification separates distinct portal vertex and penton vertex reconstructions.

**Supplementary Figure S5.**
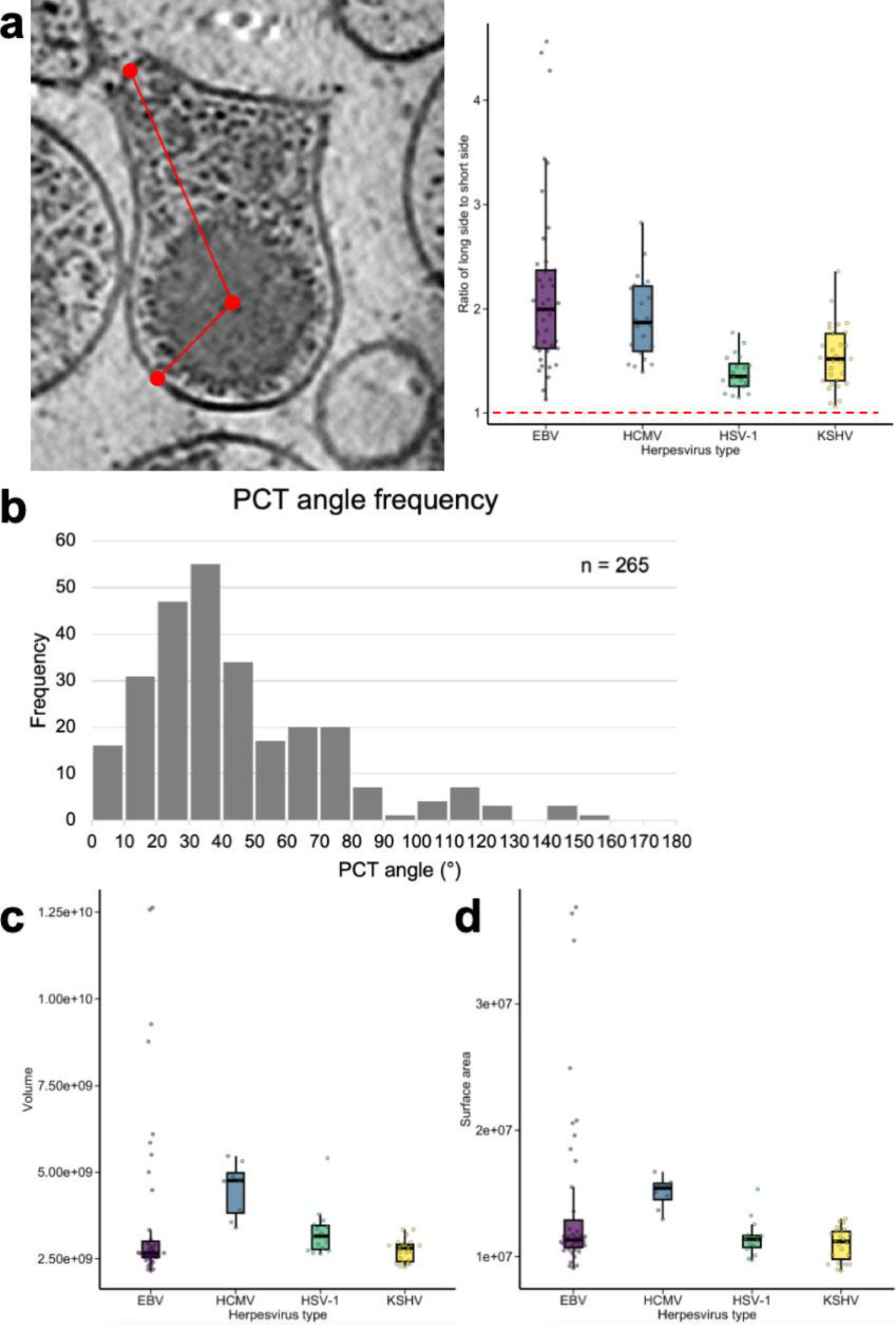
Herpesvirus subfamily morphology comparison. (a) Eccentricity of virion measured by ratio of long side to short side (left). Plot of ratios of long side to short side for virions of EBV, KSHV, HSV-1, and HCMV. Red dashed line indicates ratio for sphere for reference. (b) Histogram showing portal-capsid-tegument (PCT) angle frequency of KSHV virions. (c) Plot of volume (Å^3^) encompassed by envelope of intact virions. (d) Plot of surface area (Å^2^) of intact virion envelopes.

**Supplementary Figure S6.**
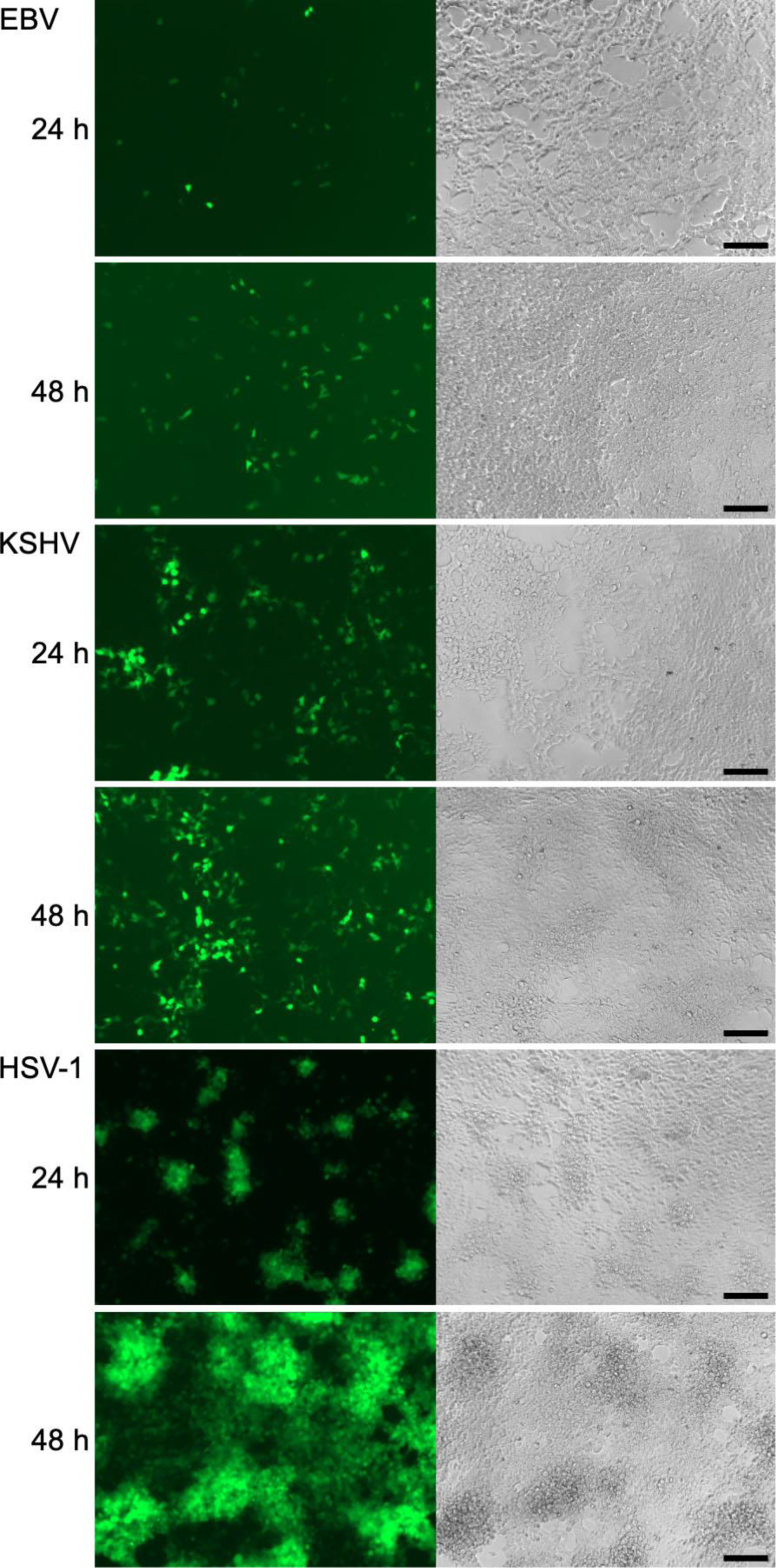
Herpesviruses retain infectivity after pelleting by ultracentrifugation. Fluorescent (left) and light (right) microscopy images of cells infected with GFP-encoding EBV, KSHV, or HSV-1 that were pelleted at 80,000 × g.

**Supplementary Figure S7.**
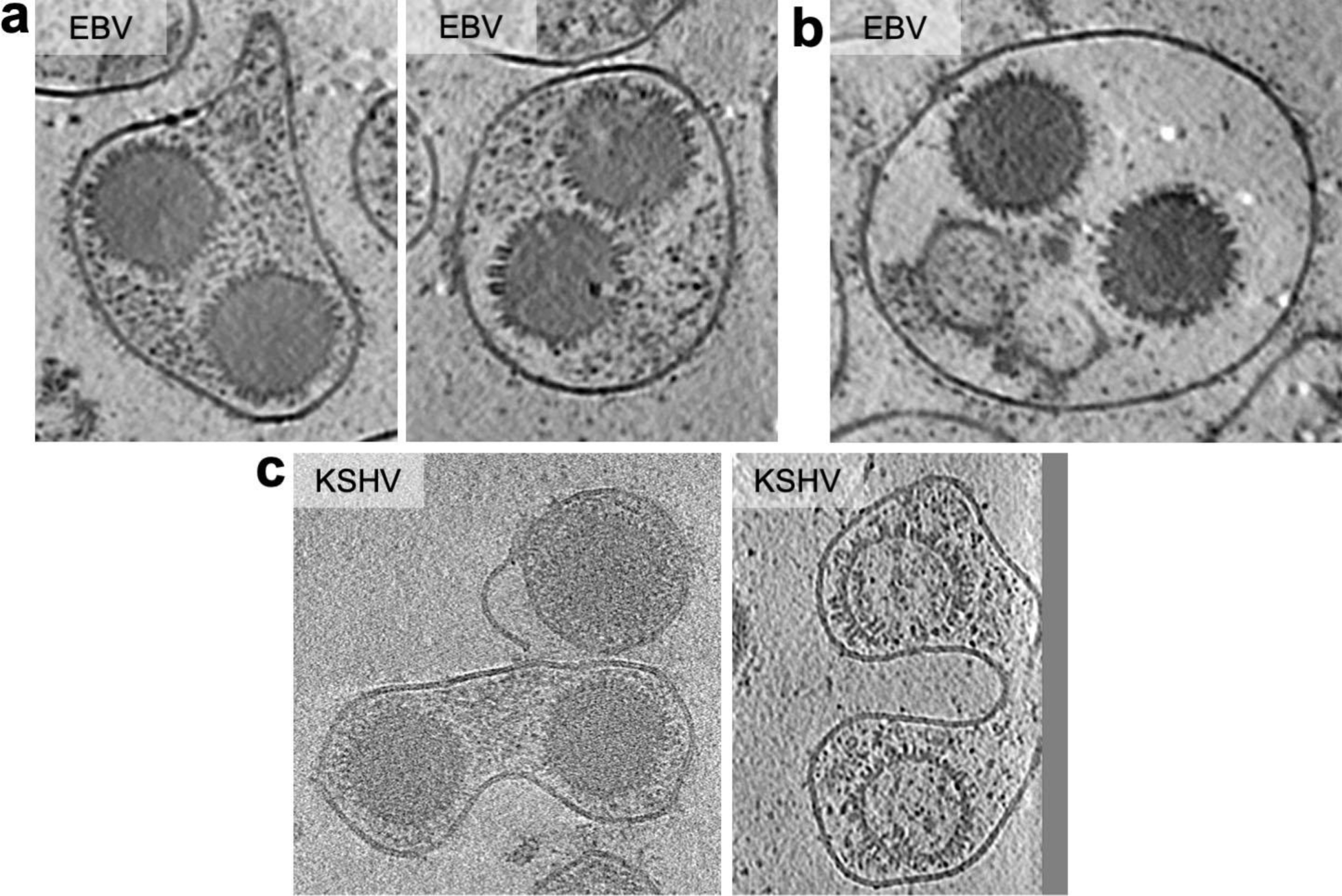
Non-standard morphologies observed in EBV and KSHV. (a) EBV particles containing two C-capsids within the same intact envelope and surrounded by tegument-like densities. (b) EBV particle containing two C-capids within an envelope with low proteinaceous density. (c) KSHV particles containing two C-capsids (left) or capsids devoid of genome (right) within the same intact envelope that is pinched inward between the capsids.

**Supplementary Table S1.**
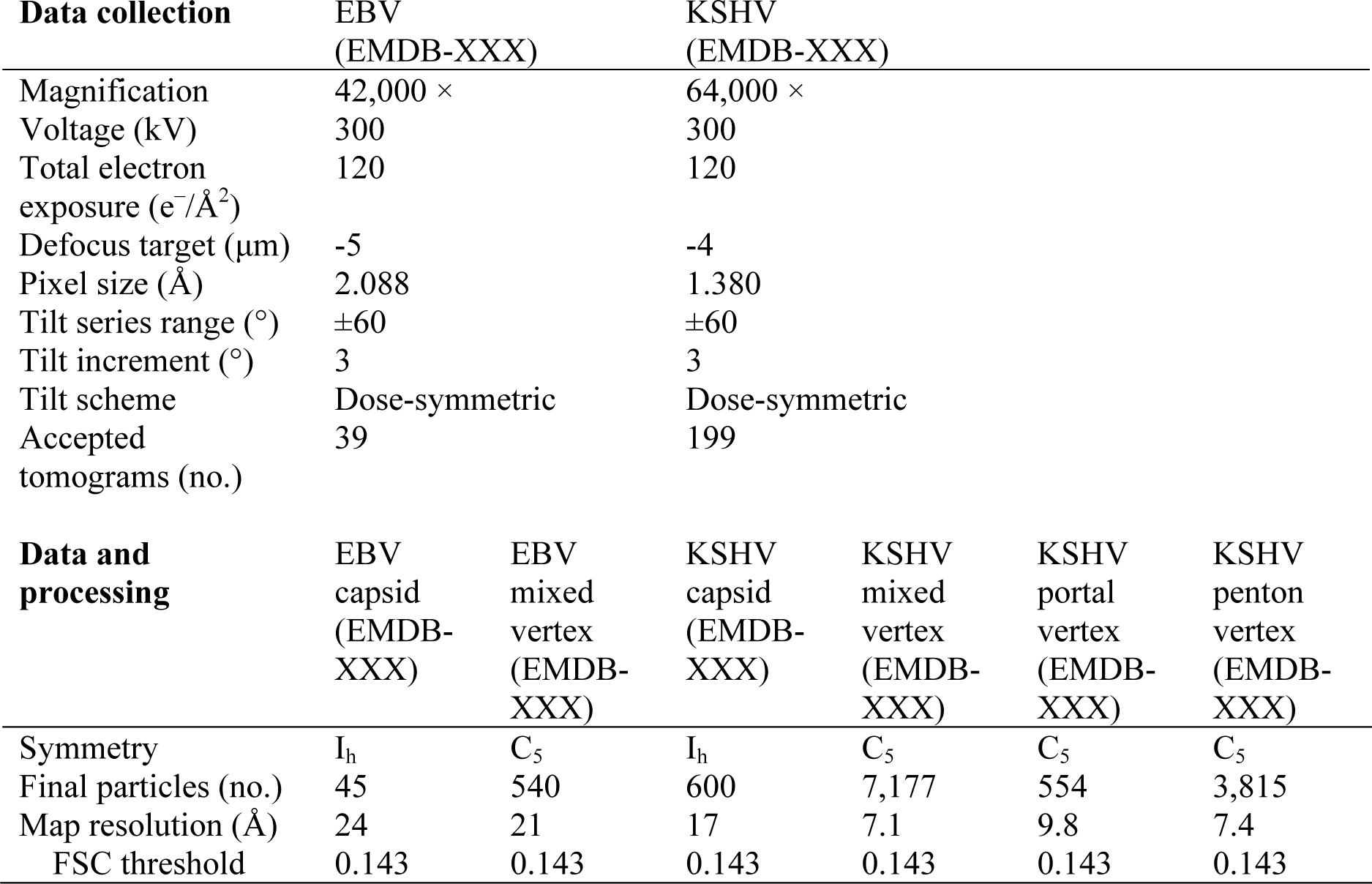
CryoET data collection and processing statistics.

| **Data collection** | EBV<br>(EMDB-XXX) |  | KSHV<br>(EMDB-XXX) |  |  |  |
| --- | --- | --- | --- | --- | --- | --- |
| Magnification | 42,000 × |  | 64,000 × |  |  |  |
| Voltage (kV) | 300 |  | 300 |  |  |  |
| Total electron exposure (e<sup>-</sup>/Å<sup>2</sup>) | 120 |  | 120 |  |  |  |
| Defocus target (μm) | -5 |  | -4 |  |  |  |
| Pixel size (Å) | 2.088 |  | 1.380 |  |  |  |
| Tilt series range (°) | ±60 |  | ±60 |  |  |  |
| Tilt increment (°) | 3 |  | 3 |  |  |  |
| Tilt scheme | Dose-symmetric |  | Dose-symmetric |  |  |  |
| Accepted tomograms (no.) | 39 |  | 199 |  |  |  |
| **Data and processing** | EBV capsid (EMDB-XXX) | EBV mixed vertex (EMDB-XXX) | KSHV capsid (EMDB-XXX) | KSHV mixed vertex (EMDB-XXX) | KSHV portal vertex (EMDB-XXX) | KSHV penton vertex (EMDB-XXX) |
| Symmetry | I<sub>h</sub> | C<sub>5</sub> | I<sub>h</sub> | C<sub>5</sub> | C<sub>5</sub> | C<sub>5</sub> |
| Final particles (no.) | 45 | 540 | 600 | 7,177 | 554 | 3,815 |
| Map resolution (Å) | 24 | 21 | 17 | 7.1 | 9.8 | 7.4 |
| FSC threshold | 0.143 | 0.143 | 0.143 | 0.143 | 0.143 | 0.143 |

## References

1. Schiller JT, Lowy DR. 2021. An Introduction to Virus Infections and Human Cancer. Recent Results Cancer Res 217:1–11.

2. Wen KW, Wang L, Menke JR, Damania B. 2022. Cancers associated with human gammaherpesviruses. The FEBS Journal 289:7631–7669.

3. Bjornevik K, Cortese M, Healy BC, Kuhle J, Mina MJ, Leng Y, Elledge SJ, Niebuhr DW, Scher AI, Munger KL, Ascherio A. 2022. Longitudinal analysis reveals high prevalence of Epstein-Barr virus associated with multiple sclerosis. Science 10.1126/science.abj8222.

4. Chang Y, Cesarman E, Pessin MS, Lee F, Culpepper J, Knowles DM, Moore PS. 1994. Identification of herpesvirus-like DNA sequences in AIDS-associated Kaposi’s sarcoma. Science 266:1865–1869.

5. Cesarman E, Damania B, Krown SE, Martin J, Bower M, Whitby D. 2019. Kaposi sarcoma. 1. Nature Reviews Disease Primers 5:1–21.

6. Cesarman E, Chang Y, Moore PS, Said JW, Knowles DM. 1995. Kaposi’s Sarcoma– Associated Herpesvirus-Like DNA Sequences in AIDS-Related Body-Cavity–Based Lymphomas. New England Journal of Medicine 332:1186–1191.

7. Soulier J, Grollet L, Oksenhendler E, Cacoub P, Cazals-Hatem D, Babinet P, d’Agay M, Clauvel J, Raphael M, Degos L, Sigaux F. 1995. Kaposi’s sarcoma-associated herpesvirus-like DNA sequences in multicentric Castleman’s disease [see comments]. Blood 86:1276–1280.

8. Davison AJ. 2007. Overview of classification, p. In Arvin, A, Campadelli-Fiume, G, Mocarski, E, Moore, PS, Roizman, B, Whitley, R, Yamanishi, K (eds.), Human Herpesviruses: Biology, Therapy, and Immunoprophylaxis. Cambridge University Press, Cambridge.

9. Cohen JI. 2020. Herpesvirus latency. J Clin Invest 130:3361–3369.

10. Mocarski Jr. ES. 2007. Comparative analysis of herpesvirus-common proteins, p. In Arvin, A, Campadelli-Fiume, G, Mocarski, E, Moore, PS, Roizman, B, Whitley, R, Yamanishi, K (eds.), Human Herpesviruses: Biology, Therapy, and Immunoprophylaxis. Cambridge University Press, Cambridge.

11. Connolly SA, Jardetzky TS, Longnecker R. 2021. The structural basis of herpesvirus entry. 2. Nature Reviews Microbiology 19:110–121.

12. Lee H-R, Lee S, Chaudhary PM, Gill P, Jung JU. 2010. Immune evasion by Kaposi’s sarcoma-associated herpesvirus. Future Microbiol 5:1349–1365.

13. Sathish N, Wang X, Yuan Y. 2012. Tegument Proteins of Kaposi’s Sarcoma-Associated Herpesvirus and Related Gamma-Herpesviruses. Frontiers in Microbiology 3.

14. Murata T. 2023. Tegument proteins of Epstein-Barr virus: Diverse functions, complex networks, and oncogenesis. Tumour Virus Research 15:200260.

15. Schrag JD, Prasad BVV, Rixon FJ, Chiu W. 1989. Three-dimensional structure of the HSV1 nucleocapsid. Cell 56:651–660.

16. Booy FP, Newcomb WW, Trus BL, Brown JC, Baker TS, Steven AC. 1991. Liquid-crystalline, phage-like packing of encapsidated DNA in herpes simplex virus. Cell 64:1007–1015.

17. Newcomb WW, Trus BL, Booy FP, Steven AC, Wall JS, Brown JC. 1993. Structure of the Herpes Simplex Virus Capsid Molecular Composition of the Pentons and the Triplexes. Journal of Molecular Biology 232:499–511.

18. Zhou ZH, Dougherty M, Jakana J, He J, Rixon FJ, Chiu W. 2000. Seeing the Herpesvirus Capsid at 8.5 Å. Science 288:877–880.

19. Steven AC, Roberts CR, Hay J, Bisher ME, Pun T, Trus BL. 1986. Hexavalent capsomers of herpes simplex virus type 2: symmetry, shape, dimensions, and oligomeric status. J Virol 57:578–584.

20. Trus BL, Heymann JB, Nealon K, Cheng N, Newcomb WW, Brown JC, Kedes DH, Steven AC. 2001. Capsid Structure of Kaposi’s Sarcoma-Associated Herpesvirus, a Gammaherpesvirus, Compared to Those of an Alphaherpesvirus, Herpes Simplex Virus Type 1, and a Betaherpesvirus, Cytomegalovirus. Journal of Virology 75:2879–2890.

21. Lo P, Yu X, Atanasov I, Chandran B, Zhou ZH. 2003. Three-Dimensional Localization of pORF65 in Kaposi’s Sarcoma-Associated Herpesvirus Capsid. Journal of Virology 77:4291–4297.

22. Yu X, Jih J, Jiang J, Zhou ZH. 2017. Atomic structure of the human cytomegalovirus capsid with its securing tegument layer of pp150. Science 356.

23. Dai X, Zhou ZH. 2018. Structure of the herpes simplex virus 1 capsid with associated tegument protein complexes. Science 360.

24. Yuan S, Wang J, Zhu D, Wang N, Gao Q, Chen W, Tang H, Wang J, Zhang X, Liu H, Rao Z, Wang X. 2018. Cryo-EM structure of a herpesvirus capsid at 3.1 Å. Science 360:eaao7283.

25. Wang J, Yuan S, Zhu D, Tang H, Wang N, Chen W, Gao Q, Li Y, Wang J, Liu H, Zhang X, Rao Z, Wang X. 2018. Structure of the herpes simplex virus type 2 C-capsid with capsid-vertex-specific component. 1. Nature Communications 9:3668.

26. Gong D, Dai X, Jih J, Liu Y-T, Bi G-Q, Sun R, Zhou ZH. 2019. DNA-Packing Portal and Capsid-Associated Tegument Complexes in the Tumor Herpesvirus KSHV. Cell 178:1329–1343.e12.

27. Liu Y-T, Jih J, Dai X, Bi G-Q, Zhou ZH. 2019. Cryo-EM structures of herpes simplex virus type 1 portal vertex and packaged genome. Nature 570:257–261.

28. Zhang Y, Liu W, Li Z, Kumar V, Alvarez-Cabrera AL, Leibovitch EC, Cui Y, Mei Y, Bi G-Q, Jacobson S, Zhou ZH. 2019. Atomic structure of the human herpesvirus 6B capsid and capsid-associated tegument complexes. 1. Nature Communications 10:5346.

29. Li Z, Zhang X, Dong L, Pang J, Xu M, Zhong Q, Zeng M-S, Yu X. 2020. CryoEM structure of the tegumented capsid of Epstein-Barr virus. Cell Research 1–12.

30. Liu W, Cui Y, Wang C, Li Z, Gong D, Dai X, Bi G-Q, Sun R, Zhou ZH. 2020. Structures of capsid and capsid-associated tegument complex inside the Epstein–Barr virus. Nature Microbiology 1–14.

31. Sun J, Liu C, Peng R, Zhang F-K, Tong Z, Liu S, Shi Y, Zhao Z, Zeng W-B, Gao GF, Shen H-J, Yang X, Luo M, Qi J, Wang P. 2020. Cryo-EM structure of the varicella-zoster virus A-capsid. 1. Nat Commun 11:4795.

32. Wang W, Zheng Q, Pan D, Yu H, Fu W, Liu J, He M, Zhu R, Cai Y, Huang Y, Zha Z, Chen Z, Ye X, Han J, Que Y, Wu T, Zhang J, Li S, Zhu H, Zhou ZH, Cheng T, Xia N. 2020. Near-atomic cryo-electron microscopy structures of varicella-zoster virus capsids. Nat Microbiol 5:1542–1552.

33. Li Z, Pang J, Dong L, Yu X. 2021. Structural basis for genome packaging, retention, and ejection in human cytomegalovirus. Nat Commun 12:4538.

34. Li Z, Pang J, Gao R, Wang Q, Zhang M, Yu X. 2023. Cryo-electron microscopy structures of capsids and in situ portals of DNA-devoid capsids of human cytomegalovirus. 1. Nat Commun 14:2025.

35. Liu Y-T, Strugatsky D, Liu W, Zhou ZH. 2021. Structure of human cytomegalovirus virion reveals host tRNA binding to capsid-associated tegument protein pp150. Nat Commun 12:5513.

36. Jih J, Liu Y-T, Liu W, Zhou ZH. 2024. The incredible bulk: Human cytomegalovirus tegument architectures uncovered by AI-empowered cryo-EM. Science Advances 10:eadj1640.

37. Cao L, Wang N, Lv Z, Chen W, Chen Z, Song L, Sha X, Wang G, Hu Y, Lian X, Cui G, Fan J, Quan Y, Liu H, Hou H, Wang X. 2024. Insights into varicella-zoster virus assembly from the B- and C-capsid at near-atomic resolution structures. hLife 2:64–74.

38. Dai X, Gong D, Lim H, Jih J, Wu T-T, Sun R, Zhou ZH. 2018. Structure and mutagenesis reveal essential capsid protein interactions for KSHV replication. Nature 553:521–525.

39. Grünewald K, Desai P, Winkler DC, Heymann JB, Belnap DM, Baumeister W, Steven AC. 2003. Three-Dimensional Structure of Herpes Simplex Virus from Cryo-Electron Tomography. Science 302:1396–1398.

40. Si Z, Zhang J, Shivakoti S, Atanasov I, Tao C-L, Hui WH, Zhou K, Yu X, Li W, Luo M, Bi G-Q, Zhou ZH. 2018. Different functional states of fusion protein gB revealed on human cytomegalovirus by cryo electron tomography with Volta phase plate. PLOS Pathogens 14:e1007452.

41. Liu Y-T, Zhang H, Wang H, Tao C-L, Bi G-Q, Zhou ZH. 2022. Isotropic reconstruction for electron tomography with deep learning. 1. Nat Commun 13:6482.

42. Deng B, O’Connor CM, Kedes DH, Zhou ZH. 2007. Direct Visualization of the Putative Portal in the Kaposi’s Sarcoma-Associated Herpesvirus Capsid by Cryoelectron Tomography. Journal of Virology 81:3640–3644.

43. Dai W, Jia Q, Bortz E, Shah S, Liu J, Atanasov I, Li X, Taylor KA, Sun R, Zhou ZH. 2008. Unique structures in a tumor herpesvirus revealed by cryo-electron tomography and microscopy. Journal of Structural Biology 161:428–438.

44. Brulois KF, Chang H, Lee AS-Y, Ensser A, Wong L-Y, Toth Z, Lee SH, Lee H-R, Myoung J, Ganem D, Oh T-K, Kim JF, Gao S-J, Jung JU. 2012. Construction and Manipulation of a New Kaposi’s Sarcoma-Associated Herpesvirus Bacterial Artificial Chromosome Clone. Journal of Virology 86:9708–9720.

45. Zivanov J, Otón J, Ke Z, von Kügelgen A, Pyle E, Qu K, Morado D, Castaño-Díez D, Zanetti G, Bharat TA, Briggs JA, Scheres SH. 2022. A Bayesian approach to single-particle electron cryo-tomography in RELION-4.0. eLife 11:e83724.

46. Gong D, Dai X, Xiao Y, Du Y, Chapa TJ, Johnson JR, Li X, Krogan NJ, Deng H, Wu T-T, Sun R. 2017. Virus-Like Vesicles of Kaposi’s Sarcoma-Associated Herpesvirus Activate Lytic Replication by Triggering Differentiation Signaling. Journal of Virology 91.

47. Baumeister W. 2005. From proteomic inventory to architecture. FEBS Letters 579:933–937.

48. Wan W, Briggs JAG. 2016. Chapter Thirteen - Cryo-Electron Tomography and Subtomogram Averaging, p. 329–367. In Crowther, RA (ed.), Methods in Enzymology. Academic Press.

49. Trus BL, Newcomb WW, Cheng N, Cardone G, Marekov L, Homa FL, Brown JC, Steven AC. 2007. Allosteric Signalling and a Nuclear Exit Strategy: Binding of UL25/UL17 Heterodimers to DNA-filled HSV-1 Capsids. Mol Cell 26:479–489.

50. Conway JF, Cockrell SK, Copeland AM, Newcomb WW, Brown JC, Homa FL. 2010. Labeling and localization of the herpes simplex virus capsid protein UL25 and its interaction with the two triplexes closest to the penton. J Mol Biol 397:575–586.

51. Toropova K, Huffman JB, Homa FL, Conway JF. 2011. The herpes simplex virus 1 UL17 protein is the second constituent of the capsid vertex-specific component required for DNA packaging and retention. J Virol 85:7513–7522.

52. Cockrell SK, Huffman JB, Toropova K, Conway JF, Homa FL. 2011. Residues of the UL25 Protein of Herpes Simplex Virus That Are Required for Its Stable Interaction with Capsids. Journal of Virology 85:4875–4887.

53. Dai X, Gong D, Wu T-T, Sun R, Zhou ZH. 2014. Organization of Capsid-Associated Tegument Components in Kaposi’s Sarcoma-Associated Herpesvirus. Journal of Virology 88:12694–12702.

54. Ilca SL, Kotecha A, Sun X, Poranen MM, Stuart DI, Huiskonen JT. 2015. Localized reconstruction of subunits from electron cryomicroscopy images of macromolecular complexes. 1. Nature Communications 6:8843.

55. Naniima P, Naimo E, Koch S, Curth U, Alkharsah KR, Ströh LJ, Binz A, Beneke J-M, Vollmer B, Böning H, Borst EM, Desai P, Bohne J, Messerle M, Bauerfeind R, Legrand P, Sodeik B, Schulz TF, Krey T. 2021. Assembly of infectious Kaposi’s sarcoma-associated herpesvirus progeny requires formation of a pORF19 pentamer. PLOS Biology 19:e3001423.

56. Bogdanow B, Gruska I, Mühlberg L, Protze J, Hohensee S, Vetter B, Bosse JB, Lehmann M, Sadeghi M, Wiebusch L, Liu F. 2023. Spatially resolved protein map of intact human cytomegalovirus virions. 9. Nat Microbiol 8:1732–1747.

57. Ibiricu I, Huiskonen JT, Döhner K, Bradke F, Sodeik B, Grünewald K. 2011. Cryo Electron Tomography of Herpes Simplex Virus during Axonal Transport and Secondary Envelopment in Primary Neurons. PLOS Pathogens 7:e1002406.

58. Liu F, Zhou ZH. 2007. Comparative virion structures of human herpesviruses, p. In Arvin, A, Campadelli-Fiume, G, Mocarski, E, Moore, PS, Roizman, B, Whitley, R, Yamanishi, K (eds.), Human Herpesviruses: Biology, Therapy, and Immunoprophylaxis. Cambridge University Press, Cambridge.

59. Baldick CJ, Shenk T. 1996. Proteins associated with purified human cytomegalovirus particles. J Virol 70:6097–6105.

60. Newcomb WW, Brown JC. 2010. Structure and Capsid Association of the Herpesvirus Large Tegument Protein UL36. Journal of Virology 84:9408–9414.

61. Fan WH, Roberts APE, McElwee M, Bhella D, Rixon FJ, Lauder R. 2014. The Large Tegument Protein pUL36 Is Essential for Formation of the Capsid Vertex-Specific Component at the Capsid-Tegument Interface of Herpes Simplex Virus 1. J Virol 89:1502–1511.

62. Liu Y-T, Shivakoti S, Jia F, Tao C-L, Zhang B, Xu F, Lau P, Bi G-Q, Zhou ZH. 2020. Biphasic exocytosis of herpesvirus from hippocampal neurons and mechanistic implication to membrane fusion. 1. Cell Discov 6:1–12.

63. Flomm FJ, Soh TK, Schneider C, Wedemann L, Britt HM, Thalassinos K, Pfitzner S, Reimer R, Grünewald K, Bosse JB. 2022. Intermittent bulk release of human cytomegalovirus. PLOS Pathogens 18:e1010575.

64. Nanbo A, Noda T, Ohba Y. 2018. Epstein–Barr Virus Acquires Its Final Envelope on Intracellular Compartments With Golgi Markers. Frontiers in Microbiology 9.

65. Chen J, Schaller S, Jardetzky TS, Longnecker R. 2020. Epstein-Barr Virus gH/gL and Kaposi’s Sarcoma-Associated Herpesvirus gH/gL Bind to Different Sites on EphA2 To Trigger Fusion. Journal of Virology 94.

66. Zeev-Ben-Mordehai T, Vasishtan D, Hernández Durán A, Vollmer B, White P, Prasad Pandurangan A, Siebert CA, Topf M, Grünewald K. 2016. Two distinct trimeric conformations of natively membrane-anchored full-length herpes simplex virus 1 glycoprotein B. Proc Natl Acad Sci U S A 113:4176–4181.

67. Vollmer B, Pražák V, Vasishtan D, Jefferys EE, Hernandez-Duran A, Vallbracht M, Klupp BG, Mettenleiter TC, Backovic M, Rey FA, Topf M, Grünewald K. 2020. The prefusion structure of herpes simplex virus glycoprotein B. Science Advances 6:eabc1726.

68. Yang S, Hiotis G, Wang Y, Chen J, Wang J, Kim M, Reinherz EL, Walz T. 2022. Dynamic HIV-1 spike motion creates vulnerability for its membrane-bound tripod to antibody attack. bioRxiv 10.1101/2022.05.09.491169.

69. Yao H, Song Y, Chen Y, Wu N, Xu J, Sun C, Zhang J, Weng T, Zhang Z, Wu Z, Cheng L, Shi D, Lu X, Lei J, Crispin M, Shi Y, Li L, Li S. 2020. Molecular Architecture of the SARS-CoV-2 Virus. Cell 183:730–738.e13.

70. Peng L, Ryazantsev S, Sun R, Zhou ZH. 2010. Three-Dimensional Visualization of Gammaherpesvirus Life Cycle in Host Cells by Electron Tomography. Structure 18:47–58.

71. Maurer UE, Sodeik B, Grünewald K. 2008. Native 3D intermediates of membrane fusion in herpes simplex virus 1 entry. Proceedings of the National Academy of Sciences 105:10559–10564.

72. Shi L, Shen Q-T, Kiel A, Wang J, Wang H-W, Melia TJ, Rothman JE, Pincet F. 2012. SNARE Proteins: One to Fuse and Three to Keep the Nascent Fusion Pore Open. Science 335:1355–1359.

73. Kim IS, Jenni S, Stanifer ML, Roth E, Whelan SPJ, van Oijen AM, Harrison SC. 2017. Mechanism of membrane fusion induced by vesicular stomatitis virus G protein. PNAS 114:E28–E36.

74. Kawabata A, Tang H, Huang H, Yamanishi K, Mori Y. 2009. Human herpesvirus 6 envelope components enriched in lipid rafts: evidence for virion-associated lipid rafts. Virol J 6:127.

75. Bender FC, Whitbeck JC, Ponce de Leon M, Lou H, Eisenberg RJ, Cohen GH. 2003. Specific Association of Glycoprotein B with Lipid Rafts during Herpes Simplex Virus Entry. Journal of Virology 77:9542–9552.

76. Lee GE, Church GA, Wilson DW. 2003. A Subpopulation of Tegument Protein vhs Localizes to Detergent-Insoluble Lipid Rafts in Herpes Simplex Virus-Infected Cells. Journal of Virology 77:2038–2045.

77. Goswami R, Shair KHY, Gershburg E. 2017. Molecular diversity of IgG responses to Epstein–Barr virus proteins in asymptomatic Epstein–Barr virus carriers. Journal of General Virology 98:2343–2350.

78. Coghill AE, Pfeiffer RM, Proietti C, Hsu W-L, Chien Y-C, Lekieffre L, Krause L, Teng A, Pablo J, Yu KJ PhD, Lou P-J MD, Wang C-P, Liu Z, Chen C-J, Middeldorp J, Mulvenna J, Bethony J, Hildesheim A, Doolan DL. 2018. Identification of a Novel, EBV-Based Antibody Risk Stratification Signature for Early Detection of Nasopharyngeal Carcinoma in Taiwan. Clinical Cancer Research 24:1305–1314.

79. Coghill AE, Fang J, Liu Z, Chen C-J, Jarrett RF, Hjalgrim H, Proietti C, Yu KJ, Hsu W-L, Lou P-J, Wang C-P, Zhao Y, Doolan DL, Hildesheim A. 2022. Identifying Epstein–Barr virus peptide sequences associated with differential IgG antibody response. International Journal of Infectious Diseases 114:65–71.

80. Labo N, Miley W, Marshall V, Gillette W, Esposito D, Bess M, Turano A, Uldrick T, Polizzotto MN, Wyvill KM, Bagni R, Yarchoan R, Whitby D. 2014. Heterogeneity and Breadth of Host Antibody Response to KSHV Infection Demonstrated by Systematic Analysis of the KSHV Proteome. PLOS Pathogens 10:e1004046.

81. Lam AK, Roshan R, Miley W, Labo N, Zhen J, Kurland AP, Cheng C, Huang H, Teng P-L, Harelson C, Gong D, Tam YK, Radu CG, Epeldegui M, Johnson JR, Zhou ZH, Whitby D, Wu T-T. 2023. Immunization of Mice with Virus-Like Vesicles of Kaposi Sarcoma-Associated Herpesvirus Reveals a Role for Antibodies Targeting ORF4 in Activating Complement-Mediated Neutralization. Journal of Virology 0:e01600–22.

82. Semmes EC, Miller IG, Wimberly CE, Phan CT, Jenks JA, Harnois MJ, Berendam SJ, Webster H, Hurst JH, Kurtzberg J, Fouda GG, Walsh KM, Permar SR. 2022. Maternal Fc-mediated non-neutralizing antibody responses correlate with protection against congenital human cytomegalovirus infection. J Clin Invest 132.

83. Greijer AE, van de Crommert JMG, Stevens SJC, Middeldorp JM. 1999. Molecular Fine-Specificity Analysis of Antibody Responses to Human Cytomegalovirus and Design of Novel Synthetic-Peptide-Based Serodiagnostic Assays. Journal of Clinical Microbiology 37:179–188.

84. Schoppel K, Kropff B, Schmidt C, Vornhagen R, Mach M. 1997. The Humoral Immune Response against Human Cytomegalovirus Is Characterized by a Delayed Synthesis of Glycoprotein-Specific Antibodies. The Journal of Infectious Diseases 175:533–544.

85. Shibamura M, Yoshikawa T, Yamada S, Inagaki T, Nguyen PHA, Fujii H, Harada S, Fukushi S, Oka A, Mizuguchi M, Saijo M. 2020. Association of human cytomegalovirus (HCMV) neutralizing antibodies with antibodies to the HCMV glycoprotein complexes. Virol J 17:120.

86. Mann DR, Hilty MD. 1982. Antibody Response to Herpes Simplex Virus Type 1 Polypeptides and Glycoproteins in Primary and Recurrent Infection. 3. Pediatr Res 16:176–180.

87. Kalantari-Dehaghi M, Chun S, Chentoufi AA, Pablo J, Liang L, Dasgupta G, Molina DM, Jasinskas A, Nakajima-Sasaki R, Felgner J, Hermanson G, BenMohamed L, Felgner PL, Davies DH. 2012. Discovery of Potential Diagnostic and Vaccine Antigens in Herpes Simplex Virus 1 and 2 by Proteome-Wide Antibody Profiling. Journal of Virology 86:4328–4339.

88. Kirchmeier M, Fluckiger A-C, Soare C, Bozic J, Ontsouka B, Ahmed T, Diress A, Pereira L, Schödel F, Plotkin S, Dalba C, Klatzmann D, Anderson DE. 2014. Enveloped Virus-Like Particle Expression of Human Cytomegalovirus Glycoprotein B Antigen Induces Antibodies with Potent and Broad Neutralizing Activity. Clinical and Vaccine Immunology 21:174–180.

89. Cairns TM, Huang Z-Y, Gallagher JR, Lin Y, Lou H, Whitbeck JC, Wald A, Cohen GH, Eisenberg RJ. 2015. Patient-Specific Neutralizing Antibody Responses to Herpes Simplex Virus Are Attributed to Epitopes on gD, gB, or Both and Can Be Type Specific. J Virol 89:9213–9231.

90. Zhang X, Hong J, Zhong L, Wu Q, Zhang S, Zhu Q, Chen H, Wei D, Li R, Zhang W, Zhang X, Wang G, Zhou X, Chen J, Kang Y, Zha Z, Duan X, Huang Y, Sun C, Kong X, Zhou Y, Chen Y, Ye X, Feng Q, Li S, Xiang T, Gao S, Zeng M-S, Zheng Q, Chen Y, Zeng Y-X, Xia N, Xu M. 2022. Protective anti-gB neutralizing antibodies targeting two vulnerable sites for EBV-cell membrane fusion. Proceedings of the National Academy of Sciences 119:e2202371119.

91. Zhong L, Zhang W, Krummenacher C, Chen Y, Zheng Q, Zhao Q, Zeng M-S, Xia N, Zeng Y-X, Xu M, Zhang X. 2023. Targeting herpesvirus entry complex and fusogen glycoproteins with prophylactic and therapeutic agents. Trends in Microbiology 31:788–804.

92. Pedro Simas J, Efstathiou S. 1998. Murine gammaherpesvirus 68: a model for the study of gammaherpesvirus pathogenesis. Trends in Microbiology 6:276–282.

93. Glaeser RM, Hall RJ. 2011. Reaching the Information Limit in Cryo-EM of Biological Macromolecules: Experimental Aspects. Biophys J 100:2331–2337.

94. Dickerson JL, Lu P-H, Hristov D, Dunin-Borkowski RE, Russo CJ. 2022. Imaging biological macromolecules in thick specimens: The role of inelastic scattering in cryoEM. Ultramicroscopy 237:113510.

95. Buijsse B, Trompenaars P, Altin V, Danev R, Glaeser RM. 2020. Spectral DQE of the Volta phase plate. Ultramicroscopy 218:113079.

96. Speck P, Longnecker R. 1999. Epstein-Barr virus (EBV) infection visualized by EGFP expression demonstrates dependence on known mediators of EBV entry. Arch Virol 144:1123–1137.

97. Dai X, Zhou ZH. 2014. Purification of Herpesvirus Virions and Capsids. Bio-protocol 4:e1193–e1193.

98. Tivol WF, Briegel A, Jensen GJ. 2008. An Improved Cryogen for Plunge Freezing. Microsc Microanal 14:375–379.

99. Hagen WJH, Wan W, Briggs JAG. 2017. Implementation of a cryo-electron tomography tilt-scheme optimized for high resolution subtomogram averaging. Journal of Structural Biology 197:191–198.

100. Mastronarde DN. 2005. Automated electron microscope tomography using robust prediction of specimen movements. Journal of Structural Biology 152:36–51.

101. Zheng SQ, Palovcak E, Armache J-P, Verba KA, Cheng Y, Agard DA. 2017. MotionCor2: anisotropic correction of beam-induced motion for improved cryo-electron microscopy. 4. Nature Methods 14:331–332.

102. Rohou A, Grigorieff N. 2015. CTFFIND4: Fast and accurate defocus estimation from electron micrographs. Journal of Structural Biology 192:216–221.

103. Mastronarde DN, Held SR. 2017. Automated tilt series alignment and tomographic reconstruction in IMOD. Journal of Structural Biology 197:102–113.

104. Heumann JM, Hoenger A, Mastronarde DN. 2011. Clustering and Variance Maps For Cryo-electron Tomography Using Wedge-Masked Differences. J Struct Biol 175:288–299.

105. Goddard TD, Huang CC, Meng EC, Pettersen EF, Couch GS, Morris JH, Ferrin TE. 2018. UCSF ChimeraX: Meeting modern challenges in visualization and analysis. Protein Science 27:14–25.

106. Ermel UH, Arghittu SM, Frangakis AS. 2022. ArtiaX: An electron tomography toolbox for the interactive handling of sub-tomograms in UCSF ChimeraX. Protein Science 31:e4472.

